# Catecholaminergic neuromodulation and selective attention jointly shape perceptual decision-making

**DOI:** 10.1101/2022.11.03.515022

**Authors:** Stijn A. Nuiten, Jan Willem De Gee, Jasper B. Zantvoord, Johannes J. Fahrenfort, Simon van Gaal

## Abstract

Perceptual decisions about sensory input are influenced by fluctuations in ongoing neural activity, most prominently driven by attention and neuromodulator systems. It is currently unknown if neuromodulator activity and attention differentially modulate perceptual decision-making and/or whether neuromodulatory systems in fact control attentional processes. To investigate the effects of two distinct neuromodulatory systems and spatial attention on perceptual decisions, we pharmacologically elevated cholinergic (through donepezil) and catecholaminergic (through atomoxetine) levels in humans performing a visuo-spatial attention task, while we measured electroencephalography (EEG). Both attention and catecholaminergic enhancement improved decision-making at the behavioral and algorithmic level, as reflected in increased perceptual sensitivity and the modulation of the drift rate parameter derived from drift diffusion modeling. Univariate analyses of EEG data time-locked to the attentional cue, the target stimulus, and the motor response, further revealed that attention and catecholaminergic enhancement both modulated pre-stimulus cortical excitability, cue- and stimulus-evoked sensory activity as well as parietal evidence accumulation signals. Interestingly, we observed both similar, unique, and interactive effects of attention and catecholaminergic neuromodulation on these behavioral, algorithmic, and neural markers of the decision-making process. Thereby, this study reveals an intricate relationship between attentional and catecholaminergic systems and advances our understanding about how these systems jointly shape various stages of perceptual decision-making.

## Introduction

Agents often make inconsistent and varying decisions when faced with identical repetitions of (sensory) evidence. Recent laboratory experiments, meticulously controlling for external factors, have shown that ongoing fluctuations in neural activity may underlie such behavioral variability (Gold & Shadlen, 2007; Renart & Machens, 2014; Waschke et al., 2021; Wyart & Koechlin, 2016). Variability in behavioral responses, however, may not be exclusively related to alterations in the decision-making process, but in fact may find its root cause in alterations in sensory processes leading up towards the decision (Benwell et al., 2017; Busch et al., 2009; Iemi et al., 2017, 2022; Samaha et al., 2020).

Candidate sources of these fluctuations in cortical activity and behavior are systematic variations in attention and central arousal state (Harris & Thiele, 2011; Summerfield & Egner, 2009). Attention to specific features of sensory input (e.g. its spatial location or content) results in facilitated processing of this input and is associated with improved behavioral performance (Carrasco, 2011; Desimone & Duncan, 1995). The central arousal state of animals is controlled by the activity of neuromodulators, including noradrenaline (NA; Aston-Jones & Cohen, 2005) and acetylcholine (ACh; Hasselmo & Sarter, 2011; McCormick, 1989), which globally innervate cortex. Contrary to attention, the relation between noradrenergic and cholinergic neuromodulator activity and behavioral performance is non-monotonic, with optimal behavioral performance occurring at intermediate levels of neuromodulation (Aston-Jones & Cohen, 2005; Bentley et al., 2011; McGinley et al., 2015). It is currently unknown if fluctuations in attention and neuromodulator activity contribute to neural and behavioral variability in the same way. Both attention and neuromodulators can alter the input/output ratio (or: gain) of single neurons and neuronal networks, thereby modulating the impact of (sensory) input on cortical processing (Aston-Jones & Cohen, 2005; Hillyard et al., 1998; Soma et al., 2012) and thus possibly perceptual experience. For example, recordings in macaque middle temporal area (MT) have provided evidence for attentional modulation of neural gain, by showing increased firing rates for neurons having their receptive field on an attended location in space, but suppressed firing rates for neurons with receptive fields outside that focus, under identical sensory input (Martinez-Trujillo & Treue, 2004). Likewise, both noradrenaline and acetylcholine are known to modulate firing rates of neurons in sensory cortices (McCormick, 1989). To illustrate, the arousal state of mice, driven by neuromodulator activity, strongly modulates ongoing membrane potentials in auditory cortex and as such can shape optimal states for auditory detection behavior (McGinley et al., 2015). Neuromodulator activity and attention also have similar effects on network dynamics, as both increase cortical desynchronization and enhance the encoding of sensory information (Harris & Thiele, 2011; Thiele & Bellgrove, 2018). These similar neural effects of attention and neuromodulators have raised the question whether neuromodulatory systems control (certain aspects of) attention (Thiele & Bellgrove, 2018).

Here, we addressed this question by investigating if and how covert spatial attention and neuromodulatory systems jointly shape, in isolation and/or in interaction, visual perception in humans. To explore how fluctuations in neuromodulatory drive affect perceptual and decision-making processes in humans, previous work has so far mainly used correlational methods, often by linking performance on simple discrimination or detection tasks to different readouts of fluctuations in brain state, most prominently variations in pupil size (de Gee et al., 2014, 2017; Murphy et al., 2014; Podvalny et al., 2021; van Kempen et al., 2019; Waschke et al., 2019; for exceptions see e.g. Beste et al., 2018; Gelbard-Sagiv et al., 2018; Loughnane et al., 2019). To provide causal evidence directly tying neuromodulatory drive to attention, we pharmacologically elevated levels of catecholamines (noradrenaline and dopamine, through atomoxetine) and acetylcholine (through donepezil) in human participants performing a probabilistic attentional cueing task, while we measured electroencephalography (EEG) and pupillary responses. Participants reported the orientation of a briefly presented Gabor patch, presented left or right of fixation, as being clockwise (CW; 45°) or counterclockwise (CCW; -45°). A visual cue predicted the location of the Gabor with 80% validity (the cue did not predict orientation, see **Figure 1A**). This set-up allowed us to test the effects of increased neuromodulator levels (drug effects) and spatial attention (cue validity effects) on several stages of cortical processing leading up to the perceptual decision about Gabor orientation. We first characterized the effects of drug condition and cue validity on perceptual sensitivity, derived from signal detection theory (Green & Swets, 1966) and latent decision parameters, derived from drift diffusion modeling (Ratcliff & McKoon, 2008). We then turned to neural data that was locked to three relevant events: cue presentation, stimulus presentation and the behavioral response. Specifically, we investigated the effects of attention and neuromodulation on accumulation of sensory evidence over time towards a decision threshold (Loughnane et al., 2019; Nieuwenhuis et al., 2005; van Kempen et al., 2019), stimulus-evoked sensory activity (Loughnane et al., 2016; Newman et al., 2017; van Kempen et al., 2019), and prestimulus anticipatory neural signals (Bauer et al., 2012; Dahl et al., 2020; Praamstra & Kourtis, 2010; Velzen & Eimer, 2003).

**Figure 1.**
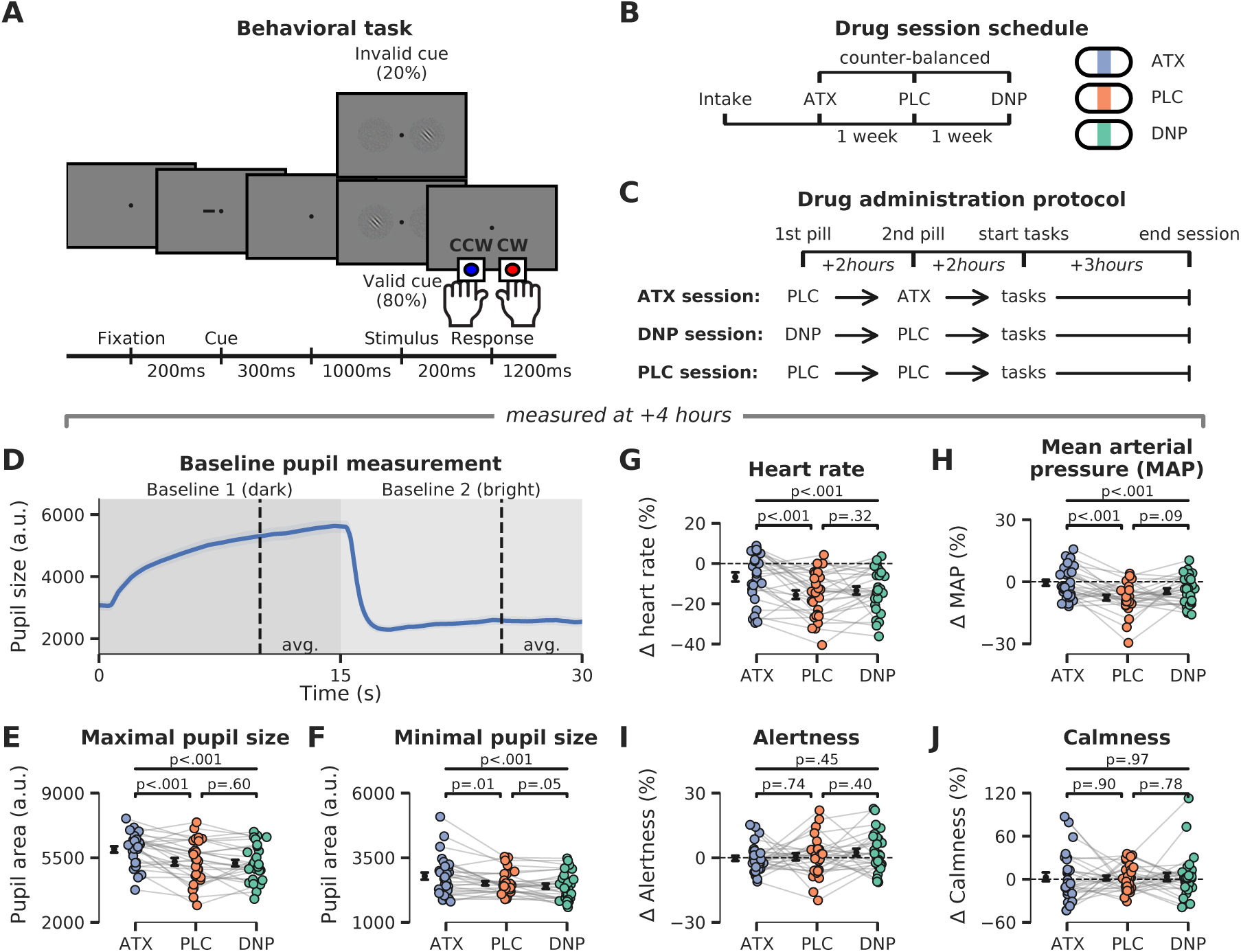
Experimental setup: behavioral task, pharmacological manipulation, and physiological responses. **A)** Schematic representation of the behavioral task. Participants responded to the orientation (CW/CCW) of unilaterally presented Gabor stimuli that were embedded in noise (bilaterally presented). The likely location of the Gabor stimulus was cued (horizontal dash presented 0.33° left/right from fixation) with 80% validity before stimulus onset. **B)** Schematic overview of experimental sessions. Participants came to the lab on four occasions: one intake session and three experimental sessions. On the experimental sessions, participants received either placebo (PLC, data in orange), donepezil (DNP, 5mg, data in green) or atomoxetine (ATX, 40mg, data in blue). Drug order was counterbalanced across participants. **C)** Time schedule of experimental sessions. Participants received a pill on two moments in each session, one at the beginning of the session and a second pill two hours later. The first pill contained either placebo (PLC and ATX session) or donepezil (DNP session), the second pill was either a placebo (PLC and DNP session) or atomoxetine (ATX session). Behavioral testing started 4 hours after administration of the first pill. **D)** Baseline pupil diameter was measured before onset of the behavioral task. Participants fixated while the background luminance of the monitor was dimmed (for 15s) and then brightened (for 15s) to establish the pupil size in dark and bright circumstances. Shading indicates standard error of the mean (SEM). **E-J)** Effects of drug on pupil diameter during the dark (P_max_, **E**) and bright (P_min_, **F**) measurement windows, heart rate (HR, **G**), mean arterial blood pressure (MAP, **H**, see **Methods**), and subjective ratings (Visual Analogue Scale, VAS; see **Methods)** of alertness (panel **I**) and calmness (**J**). All measurements except pupil diameter were baseline-corrected to the first measurement taken right before ingestion of the first pill.

## Results

### Catecholaminergic enhancement increases physiological markers of bodily arousal

We employed a randomized, double-blind crossover design in which atomoxetine (ATX, 40mg), donepezil (DNP, 5mg) and placebo (PLC) were administered in different EEG recording sessions (drug order was counterbalanced between participants, **Figure 1B**). ATX is a relatively selective noradrenaline reuptake inhibitor, which inhibits the presynaptic noradrenaline reuptake transporter, thereby resulting in increased NA and dopamine (DA) levels (Simpson & Plosker, 2004). DNP is a cholinesterase inhibitor, which impedes the breakdown of ACh by cholinesterase, thereby resulting in overall increased ACh levels (Rogers & Friedhoff, 1998). Our drug administration procedure was set up to take the difference in time to reach maximum plasma concentrations (T_max_) of DNP (∼four hours) and ATX (∼two hours) into account, while keeping participants blind to the current drug condition. Because T_max_ was different for ATX and DNP and we did not want to introduce any differences in terms of the pill-taking regime, we added a placebo (PLC) pill within all the experimental sessions, also for the ATX and the DNP protocol. Thus, participants would always ingest pills at the same moments in time, regardless of the session they were in. On every experimental session, we always administered two pills at the same moments in time, differing only in which of these two pills contained the drug. Specifically, in the DNP session this meant that the first pill was DNP followed two hours later by PLC, while for the ATX session this meant that the first pill was PLC followed two hours later by ATX. In the PLC session both pills were PLC (see **Figure 1C** below).

To gauge the effect of our pharmaceuticals, we collected physiological and subjective state measures at different moments throughout the day (from 9:00 – 16:00; **Figure 1D-1J**; for details see **Methods**). We performed one-way repeated measures (rm)ANOVAs on these physiological and subjective measures, to test for omnibus main effects of drug condition. Next, we used post-hoc paired-sample t-tests to test for pairwise drug effects (ATX/DNP vs. PLC). Throughout this work, all effect sizes are expressed as ƞ^2^_*p*_ and should be interpreted as medium sized for ƞ^2^_*p*_>0.06 and large for ƞ^2^_*p*_>0.14 (Cohen, 1988).

Prior to onset of the behavioral tasks, we observed strong effects of drug condition on pupil size (P_max_: F_2,54_=15.84, p<.001, ƞ^2^_*p*_=0.37; P_min_: F_2,54_=10.56, p<.001, ƞ^2^_*p*_=0.28), heart rate (F_2,54_=8.36, p<.001, ƞ^2^_*p*_=0.24), and ƞ^2^_*p*_ mean arterial blood pressure (F_2,54_=8.05, p<.001, ƞ^2^_*p*_=0.23). Post-hoc tests indicated that all physiological measures were only affected by ATX (P_max_: t(27)=4.12, p<.001, ƞ^2^_*p*_=0.39; P_min_: t(27)=2.77, p=.01, ƞ^2^_*p*_=0.22; HR: t(27)=4.11, p<.001, ƞ^2^_*p*_=0.38; MAP: t(27)=4.33, p<.001, ƞ^2^_*p*_=0.41) and not DNP (P_max_: t(27)=-0.52, p=.60, ƞ^2^_*p*_=0.01, BF_01_=4.11; P_min_: t(27)=-2.02, p=.05, ƞ^2^_*p*_=0.13, note this is a trend in the opposite direction; HR: t(27)=1.01, p=.32, ƞ^2^_*p*_=0.04, BF_01_=3.13; MAP: t(27)=1.76, p=.09, ƞ^2^_*p*_=0.10, BF_01_=0.78). The robust effects of ATX, but absence of effects under DNP are in line with previous non-clinical reports (Pfeffer et al., 2018, 2021). Moreover, the effects of ATX on physiological measures of arousal were long lasting (**Figure 1 – Supplement 1**). We performed an additional multivariate ANOVA to test whether grouping the physiological responses (HR, MAP and pupil) would elucidate global effects of DNP, but this was not the case (Wilks’ lambda=0.93, F_5,50_=0.99, p=.42). Drug condition did not modulate subjective measures of alertness (F_2,54_=0.82, p=.45, ƞ^2^_*p*_=0.03) and calmness (F_2,54_=0.03, p=.97, ƞ^2^_*p*_=0.00), suggesting that participants were not aware of their heightened arousal state. However, forced-choice guessing at the end of the day about receiving any active substance (ATX, DNP) or not (PLC) suggested that some participants may have been aware of being in a drug session (proportion z-tests ATX vs. PLC: z=2.96, p=.003; DNP vs. PLC: z=0.55, p=.10).

### Catecholaminergic enhancement and attention modulate the rate of sensory evidence accumulation

To establish if and how cue validity and drug condition affected perceptual decision-making, we tested their respective effects on both the outcomes and latent constituents of the decision-making process. First, we report the effects of cue validity and drug on outcome measures of the decision, quantified with perceptual sensitivity (d’), derived from Signal Detection Theory (SDT; Green & Swets, 1966), and reaction times (RT). Specifically, first we used 3x2 factor (drug x cue validity) repeated measures (rm)ANOVAs to test for omnibus effects of cue validity, drug condition and their interactions (leveraging the full scope of data in our design). Thereafter, we performed planned 2x2 factor (specific drug x cue validity) rmANOVAs to test for pairwise main and interaction effects of each active drug (ATX/DNP vs. PLC, see **Methods** for detailed description of statistical analyses). These drug specific ANOVAs test for drug modulations that may be obscured in the full factor ANOVA.

As expected, validly cued (attended) targets were associated with improved d’ and increased response speed compared to invalidly cued (unattended) targets (main effects, d’: F_1,27_=39.22, p<.001, ƞ^2^_*p*_=0.59; RT: F_1,27_=43.47, p<.001, ƞ^2^_*p*_=0.62). This beneficial effect of cue validity on d’ was not modulated by drug (F_2,54_=2.11, p=.13, ƞ^2^_*p*_=0.07. BF_01_=1.87; **Figure 2B**), and was present in both drug conditions, although post-hoc tests showed a trending interaction between cue validity and ATX vs PLC (F_1,27_=3.76, p=.06, ƞ^2^_*p*_=0.12), but not for DNP vs PLC (F_1,27_=1.61, p=.22, ƞ^2^_*p*_=0.06, BF_01_=2.00). The beneficial effect of cue validity on RT was not modulated by drug (F_2,54_=1.35, p=.27, ƞ^2^_*p*_=0.05, BF_01_=3.33). The effect of drug condition on behavior was more subtle compared to the effects of cue validity. First, the main effect of drug condition on d’ was not significant (F_2,54_=2.80, p=.07, ƞ^2^_*p*_=0.09; **Figure 2A**), but planned pairwise rmANOVAs indicated that d’ was improved by ATX, but not by DNP, compared to PLC (ATX: F_1,27_=4.28, p=.048, ƞ^2^_*p*_=0.14; DNP: F_1,27_=1.61, p=.22, ƞ^2^_*p*_=0.06, BF_01_=1.58; **Figure 2A**). RTs were not affected by drug (F_2,54_=1.33, p=.27, ƞ^2^_*p*_=0.05, BF_01_=1.99; **Figure 2B**), and pairwise comparisons (vs. PLC) were also not significant for neither ATX nor DNP (ATX: F_1,27_=1.39, p=.25, ƞ^2^_*p*_=0.05, BF_01_=1.59; DNP: F_1,27_=2.82, p=.11, ƞ^2^_*p*_=0.10, BF_01_=0.95). Participants did not exhibit changes in response bias for the different condition (for the full results see **Figure 2 - Supplement 1**).

**Figure 2.**
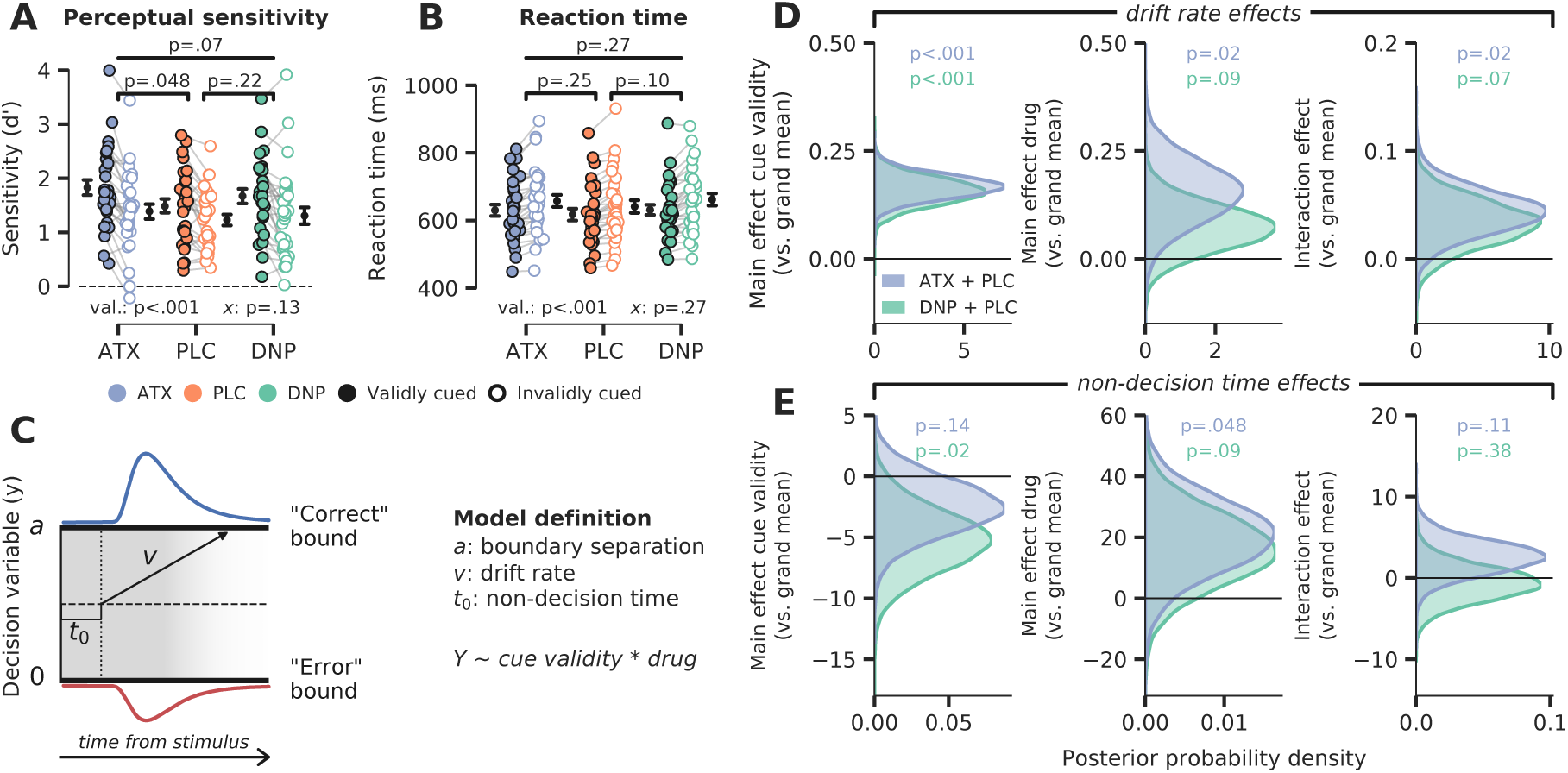
Behavioral results. **A)** Signal detection theoretic sensitivity (d’), separately per drug and cue validity. Error bars indicate SEM, *x* demarks the p-value for the omnibus interaction effect between drug condition and cue validity, val. refers to the factor cue validity **B)** As A, but for reaction time (RT). **C)** Schematic of the drift diffusion model (DDM), accounting for behavioral performance and reaction times. The model describes behavior based on various latent parameters, including drift rate (*v*), boundary separation (*a*) and non-decision time (*t_0_*). These parameters (demarked with *Y* in formula) were allowed to fluctuate with cue validity, drug condition and their interaction. Models were fitted separately for ATX (+ PLC) and DNP (+ PLC). **D-E)** Posterior probability distributions for DDM parameter estimates (blue: ATX model, green: DNP model). The effects of cue validity (left column), drug (middle column) and their interaction (right column) are shown for **D)** drift rate and **E)** non-decision time.

Perceptual sensitivity and RT are behavioral measures that only capture the final output of the decision process and are not directly informative with respect to the components underlying the formation of a decision. Computational models that are fitted to behavioral data fill this gap by decomposing choice behavior into constituent cognitive processes (Forstmann et al., 2016). Besides this, another advantage of computational modelling over analyses of choice behavior is that it integrates various dependent variables -i.e. choice accuracy and response time distributions-resulting in more robust descriptions of the underlying data. Here, we aimed to quantify the effects of drug and cue validity on latent variables of the decision-making process, by fitting a drift-diffusion model (DDM; Ratcliff & McKoon, 2008) to behavioral data (see **Methods**). The DDM describes perceptual decisions as the accumulation of noisy evidence over time that is terminated when accumulated evidence crosses a certain decision threshold (**Figure 2C**). The basic version of the model consists of three variables: drift rate (reflecting the average rate of evidence accumulation; *v*), decision boundary separation (reflecting the amount of evidence required to commit to a decision; *b*), and non-decision time (the duration of non-decision processes, including sensory encoding and motor execution; *t0*). Based on previous literature, we hypothesized that cue validity would mainly affect sensory evidence accumulation reflected in drift rate (Loughnane et al., 2016; van Vugt et al., 2019), further suggested by the fact that valid cues resulted in better and faster performance on Gabor discrimination (**Figure 2A, B**). On the contrary, although perceptual sensitivity improved under ATX (vs. PLC), average RTs were not affected, suggesting that other parameters may also have been modulated by drug besides drift rate. So, to tease apart the effects of cue validity and drug condition, we fitted a model in which drift rate, boundary separation and non-decision time were allowed to fluctuate with cue validity and drug. We included drift rate variability to account for between trial variance in drift rate. We fitted two hierarchical Bayesian regression models to capture within-subject effects of both drug and cue validity: one for ATX vs. PLC and one for DNP vs. PLC. Note that these regression models did not allow to include all three drug conditions in one model and therefore we applied a 2x2 factorial design (cue validity x drug) for both ATX and DNP separately. The Bayesian implementation of the DDM constrained single-subject parameter estimates based on group-level parameter distributions, thereby increasing robustness to outliers. Model fits are reported in **Figure 2 – Supplement 2**.

We indeed found that validly cued trials were associated with increased drift rates. This can be observed in **Figure 2D** by a clear divergence of the posterior probability distributions for ATX vs PLC (in blue) and DNP vs PLC (in green) from 0 (both models p<.001; **Figure 2D**, “cue validity effects”). We also observed increased drift rate under ATX (in blue), but not for DNP (ATX model: p=.02, DNP model: p=.09; **Figure 2D** “drug effects”). The means of the posterior distributions (an indication of effect size) for the effects of cue validity and ATX on drift rate were comparable (ATX model: x̄ _cue_=0.17, x̄ _ATX_=0.15), although the estimates of cue validity effects were more precise than those of ATX (ATX model: s_cue_=0.03, s_ATX_=0.07). Moreover, ATX increased the effects of cue validity on drift rate, but the effect size was small in comparison to the main effects of ATX and cue validity (ATX model: p=.02; x̄ _interaction_=0.04; **Figure 2D**, “interaction effects”). DNP did not significantly modulate the effects of cue validity on drift rate, although the direction of this modulation was similar to ATX (DNP model: p=.07; **Figure 2D**).

Although the rate of evidence accumulation was increased under ATX, the onset of this accumulation was delayed, indexed by an increase in non-decision time (ATX model: p=.048, DNP model: p=.09; **Figure 2E**). The effect of cue validity on non-decision time was not robust, when taking both models into account, although the effect was significant in the DNP model (ATX model: p=.14, DNP model: p=.02; **Figure 2E**). We also did not observe any interaction effects between cue validity and drug on non-decision time (ATX model: p=.11, DNP model: p=.38; **Figure 2E**). Cue validity and drug did not affect decision bound separation, showing that response caution was not under the influence of neuromodulation or attention (**Figure 2 – Supplement 3**). We fitted two additional models (one for ATX vs. PLC and one for DNP vs. PLC) that allowed drift rate variability to also fluctuate with cue validity and drug, while keeping decision bound fixed. We used this model to test whether these effects were indeed related to overall drift rate and not drift rate variability (Murphy et al., 2014). We observed no effects of cue validity or drug on drift rate variability showing that the overall rate, and not the consistency, of evidence accumulation was enhanced by ATX and cue validity (**Figure 2 – Supplement 4**).

In sum, the DDM results revealed robust effects of attention and catecholaminergic enhancement on decision-making at the algorithmic level. Valid cues and ATX both enhanced the rate of evidence accumulation towards a decision threshold and ATX also increased the effect of cue validity on drift rate, although this interaction effect was smaller than the respective main effects of ATX and cue validity. Cholinergic main and interaction effects were often in the same direction as for ATX, but not statistically reliable. These DDM results were echoed in the behavioral performance measures (especially d’), although less robustly, highlighting the benefits of combining several behavioral measures (accuracy and RT) into drift diffusion modelling.

### Neuromodulatory changes in evidence accumulation reflected in centro-parietal accumulation signal

Evidence accumulation (defined at the algorithmic level as drift rate) is thought to be reflected in neural activity in centro-parietal cortex (Gold & Shadlen, 2007). A recently identified positively trending signal over human centro-parietal regions, termed the centro-parietal positivity (CPP), has been shown to track task difficulty, RTs, and task performance during challenging perceptual decisions (O’Connell et al., 2012; van Vugt et al., 2019). The CPP furthermore scales independently of stimulus identity (unsigned component, identical for two or more stimulus categories), correlates with drift rate as derived from modelling behavior with DDMs and as such is regarded as a neural reflection of the decision variable (i.e. accumulated evidence, **Figure 2C**; Twomey et al., 2015).

We plotted the CPP locked to the response in **Figure 3**. Note that we have transformed the EEG data to current source density (CSD) to sharpen the spatial sensitivity of our EEG results, which is suggested to be optimal for CPP analyses (Kelly & O’Connell, 2013; non-CSD transformed topographic maps can be observed in **Figure 3 – Supplement 1**). In line with previous studies (O’Connell et al., 2012; van Vugt et al., 2019), the CPP predicted the accuracy and speed of perceptual decisions: CPP amplitude at the time of the response and its slope in the time-window starting -250ms preceding the response were both higher for correct than erroneous decisions (amplitude: F_1,27_=34.28, p<.001, ƞ^2^_*p*_=0.56; slope: F_1,27_=33.20, p<.001, ƞ^2^_*p*_=0.55; **Figure 3A**), whereas decision speed was uniquely associated with CPP slope (amplitude: F_1,27_=2.06, p=.16, ƞ^2^_*p*_=0.07, BF_01_=1.60; slope: F_1,27_=10.23, p=.004, ƞ^2^_*p*_=0.27; **Figure 3B**). Moreover, subjects with high drift rate (across all conditions, median split) showed both higher CPP amplitude and slope (one-way ANOVA; amplitude: F_1,26_=6.83, p=.01, ƞ^2^_*p*_=0.21; slope: F_1,26_=5.03, p=.03, ƞ^2^_*p*_=0.16; **Figure 3C**), indicating that the CPP indeed reflects accumulated evidence over time (O’Connell et al., 2012; Twomey et al., 2015).

**Figure 3.**
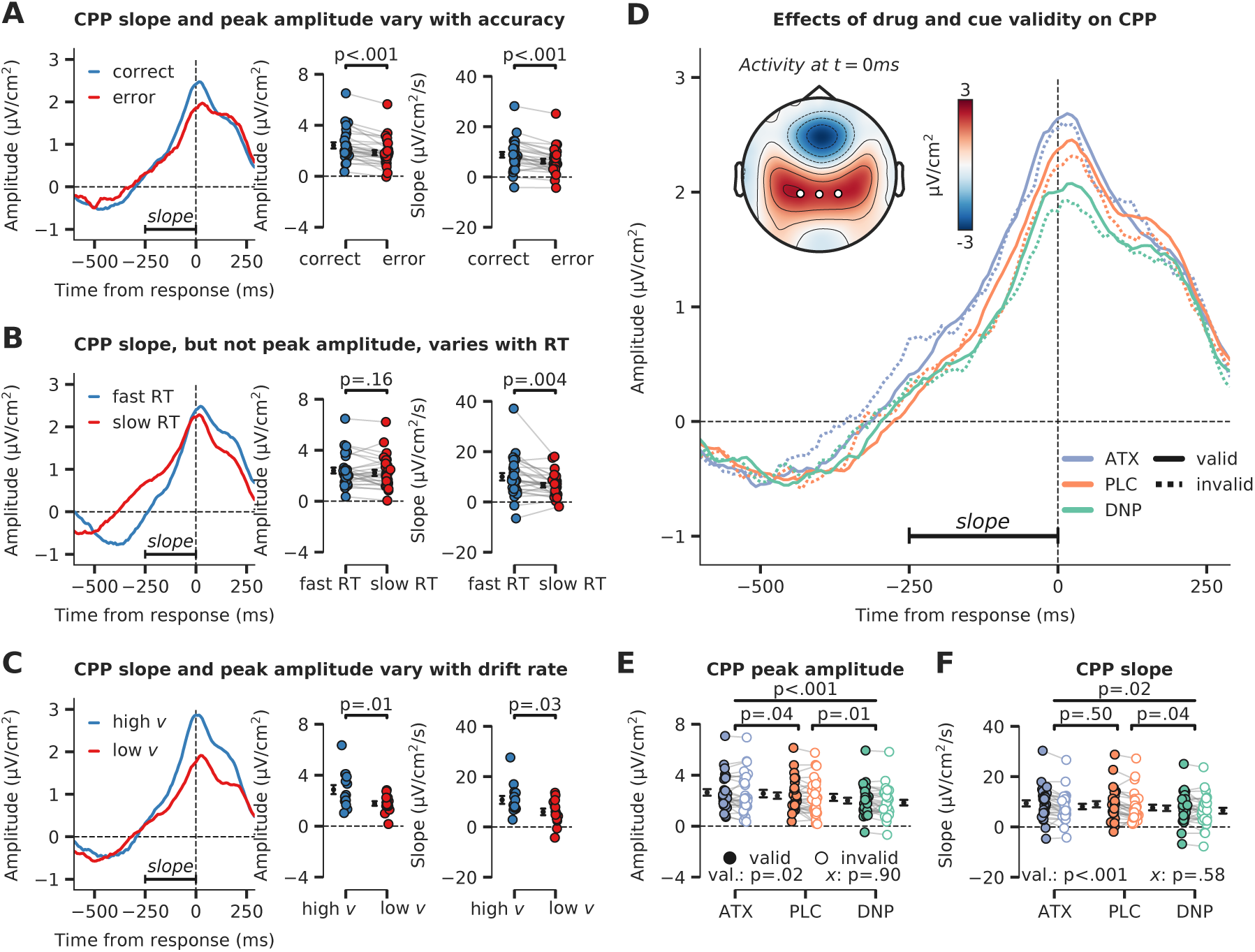
Evidence accumulation is affected by cue validity and drug, indexed by changes in centroparietal positivity (CPP). **A)** Response-locked CPP for correct and incorrect answers, **B)** for trials with fast and slow RTs and **C)** for participants with overall high drift rate and low drift rate. **D)** Modulation of response-locked CPP by drug and cue validity. The horizontal black line indicates the time-window for which CPP slope was calculated (linear regression from -250ms to 0ms pre-response). The topographic map shows activation at the moment of the response, with white markers indicating the centro-parietal ROI used for the CPP analyses (channels CP1, CP2, CPz). **E)** Peak CPP amplitude, separately for drug and cue validity. **F)** CPP slope for all drug and cue validity conditions. Note that *x* demarks the p-value for the omnibus interaction effect between drug condition and cue validity and that val. (short for validity) refers to the factor cue validity. Error bars indicate SEM.

After establishing the CPP as a reliable marker of evidence accumulation, we inspected its relation to neuromodulatory drive and cue validity (**Figure 3D**). Interestingly, drug condition affected the amplitude and slope of the CPP in a robust manner (main effects of drug, amplitude: F_2,54_=11.26, p<.001, ƞ^2^_*p*_=0.29, **Figure 3E**; slope: F_2,54_=4.31, p=.02, ƞ^2^_*p*_=0.14, **Figure 3F**). Specifically, CPP amplitude was heightened by ATX compared to PLC (amplitude: F_1,27_=4.73, p=.04, ƞ^2^_*p*_=0.15; slope: F_1,27_=0.47, p=.50, ƞ^2^_*p*_=0.02, BF_01_=4.08), whereas DNP lowered CPP peak amplitude and decreased its slope compared to PLC (amplitude: F_1,27_=8.71, p=.01, ƞ^2^_*p*_=0.24; slope: F_1,27_=4.91, p=.04, ƞ^2^_*p*_=0.15). Trials on which target locations were validly cued were associated with an increased CPP slope and amplitude (amplitude: F_1,27_=6.13, p=.02, ƞ^2^_*p*_=0.19, **Figure 3E**; slope: F_1,27_=14.70, p<.001, ƞ^2^_*p*_=0.35, **Figure 3F**). Drug condition and cue validity shaped CPP slope and peak amplitude independently because no significant interactions were observed (amplitude: F_2,54_=0.10, p=.90, ƞ^2^_*p*_=0.00, BF_01_=8.53, **Figure 3E**; slope: F_2,54_=0.55, p=.58, ƞ^2^_*p*_=0.02, BF_01_=6.08, **Figure 3F**, Bayesian evidence was in favor of the null). Summarizing, we show that drug condition and cue validity both affected CPP peak amplitude (ATX, DNP and cue validity) and slope (DNP and cue validity) and interactions between these factors were not observed.

### Stimulus-locked signals over occipito-temporal sites precede evidence accumulation and are modulated by catecholaminergic stimulation and attention

Previous studies investigating the neural origins of perceptual decision-making during visual tasks have observed early EEG deflections at lateral occipito-temporal sites that reflect target selection prior to CPP build-up (Loughnane et al., 2016; Newman et al., 2017; van Kempen et al., 2019). These target related signals in perceptual decision tasks are usually measured at electrode sites P7/P9 (left hemisphere) and P8/P10 (right hemisphere) and are referred to as the N2c (contralateral to the target) and N2i (ipsilateral to the target; (Loughnane et al., 2016; van Kempen et al., 2019). Similar to the CPP, N2c/i amplitudes scale with RT and stimulus coherence and shifts in the peak latency of the N2c have been shown to influence the onset of the CPP (Loughnane et al., 2016), suggesting that these signals may strongly modulate subsequent evidence accumulation signals.

To test whether the effects of increased neuromodulation and attention on the CPP were preceded by modulations of these occipito-temporal signals, we extracted stimulus-locked neural activity from these posterior lateral electrodes (based on literature; Loughnane et al., 2016; Newman et al., 2017; Papaioannou & Luck, 2020; van Kempen et al., 2019) as spatial regions of interest (left ROI: P7/P9; right ROI: P8/P10; **Figure 4A**). Next, we performed a cluster-corrected 2x2x3 factorial rmANOVA across time to test for effects of the cue (cued vs uncued), stimulus identity (target/noise) and drug (ATX/DNP/PLC), as well as possible interaction effects. To avoid potential confusion, the relationship between the factors cue and stimulus identity is further explicated in **Figure 4A**. We plotted the ERP traces for all 12 conditions in **Figure 4B** (stimulus-locked). To anticipate the results, we only observed main effects of each of the three factors and no interactions. In **Figure 4C**, we plotted the main effects of cue (left panel, stimulus cued vs. stimulus uncued), stimulus identity (middle panel, target vs. noise stimulus) and drug condition (right panel, all three drugs). For each main effect plot, data were averaged over the other experimental factors. Below we report the specific effects in more detail.

**Figure 4.**
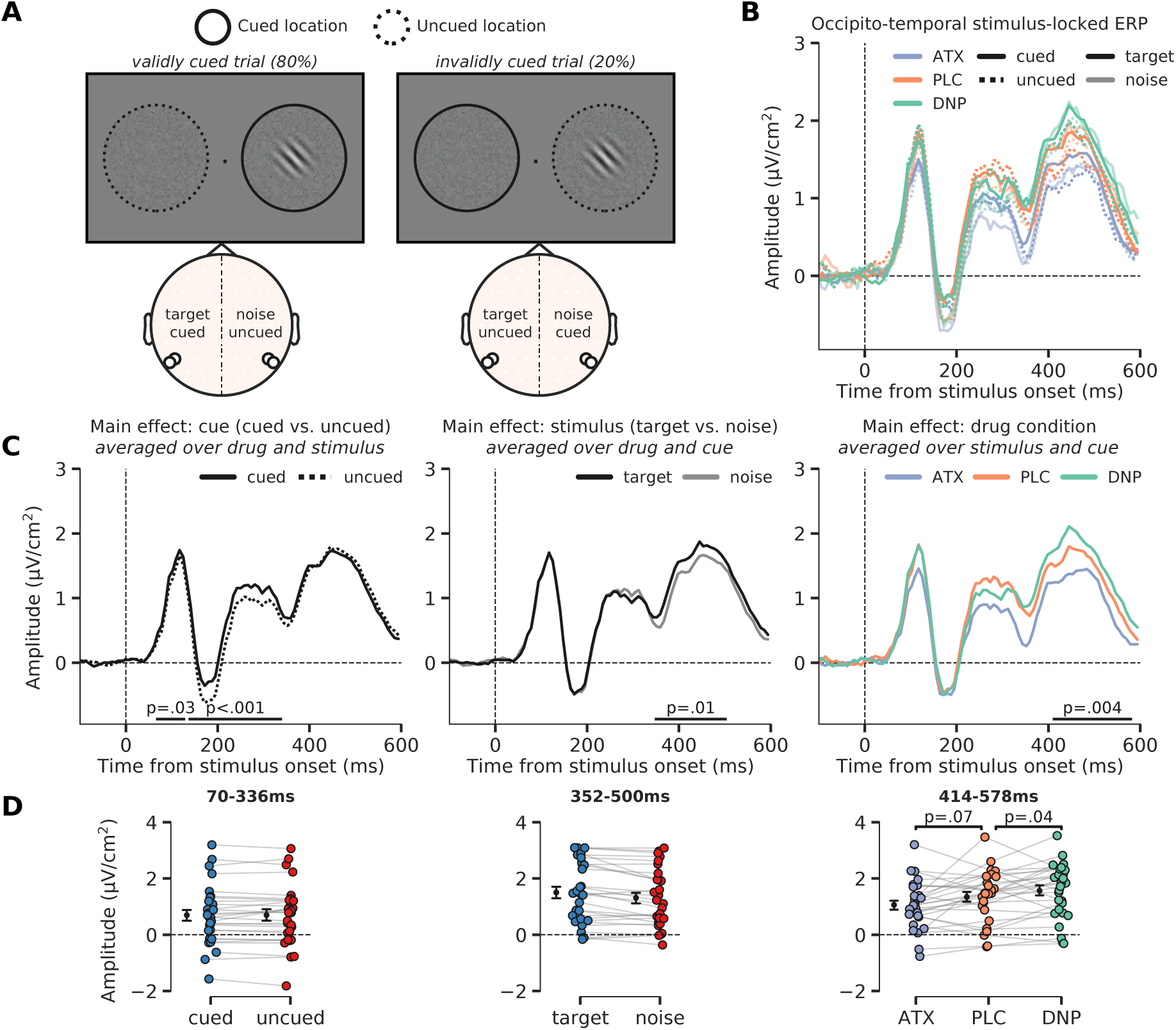
Modulation of perceptual processing by drug and attention, indexed by changes in occipito-temporal activity. **A**) Schematic of the definition of ERP conditions, based on the cued location and the location of the target stimulus. The cue was presented before the stimulus and is not shown here, but the cued and uncued location are, respectively, illustrated by solid and dashed circles around the stimulus (not shown in the actual experiment). We extracted activity from bilateral occipito-temporal ROIs (left ROI: P7/P9 and right ROI: P8/P10; see white markers in topographic map), separate for target stimuli (ROI contralateral to target) and noise stimuli (ROI ipsilateral to the target) and cued stimuli (ROI contralateral to cued location) and uncued stimuli (ROI ipsilateral to cued location). This was done for each drug condition. In the schematic we show two example scenarios for when the target stimulus appeared on the right side of fixation (it could also appear on the left side of fixation). In the first example, in the left panel, the target stimulus was cued (validly cued trial), in which case the left ROI shows activity related to the ‘cued target’ and the right ROI shows activity related to the ‘uncued noise’ stimulus. In the second example, in the right panel, the noise stimulus was cued (invalidly cued trial) in which case the right ROI shows activity related to the ‘cued noise’ stimulus and the left ROI shows activity related to the ‘uncued target’ stimulus. By defining ERP traces as such, we were able to investigate the effects of the spatial cue in isolation (i.e. cued vs. uncued, spatial attention effect) and in interaction with stimulus identity (i.e. cue validity effect). **B)** Stimulus-locked ERPs (0ms = stimulus presentation) over regions processing the target stimulus (i.e. contralateral to target stimulus, opaque line) and the noise stimulus (i.e. ipsilateral to target stimulus and/or contralateral to noise stimulus, transparent line). These traces are further split up depending on cue condition, meaning whether the stimulus was cued (i.e. contralateral to the cue, continuous line) or uncued (i.e. ipsilateral to the cue, dashed line). Note that the traces of *validly cued* trials can be seen in both cued target (solid, opaque lines) and uncued noise trials (dashed, transparent lines). Finally, the traces are plotted for the different drug conditions. **C)** The main effects of cue (cued vs uncued, left panel), stimulus identity (target vs. noise, middle panel) and drug condition (ATC/DNP/PLC, right panel). **D)** Data extracted from these cluster for each of the conditions of the main effect. Post-hoc two-sides t-test were performed to test the respective effects of ATX and DNP vs. PLC (right panel).

First, cued stimuli triggered a more positive (or less negative) ERP response than uncued stimuli already early on, specifically from 70ms-336ms post-stimulus (two clusters: 70ms-125ms, p=.043; 141ms-336ms, p<.001; **Figure 4C – left panel**). The modulation of this activity (averaged across both temporal clusters: 70-336ms) was similar for when the target stimulus was cued vs. when the noise stimulus was cued and therefore reflects an initial modulation of visual processing of any cued (attended) stimulus, irrespective of whether it was a target or not (F_1,27_=0.08, p=.77, ƞ^2^_*p*_=0.00, BF_01_=4.58; **Figure 4 – Supplement 1A**). In other words, there was no cue validity effect, which would have been reflected in an interaction between cue and stimulus identity. Moreover, the cue related modulation of neural activity (cued vs. uncued) was not affected by drug condition (F_2,54_=0.01, p=.99, ƞ^2^_*p*_=0.00, BF_01_=8.21) and we also observed no 3-way interaction between cue, stimulus and drug in this time-window (F_2,54_=0.57, p=.57, ƞ^2^_*p*_=0.02, BF_01_=3.76; **Figure 4 – Supplement 1A**).

Second, the main effect of stimulus identity on occipito-temporal activity (difference between target vs. noise stimuli) occurred later in time, from 352-500ms post-stimulus (p=.01; **Figure 4C – middle panel**). This effect was not modulated by drug condition (F_2,54_=0.74, p=.48, ƞ^2^_*p*_=0.03, BF_01_=9.63) or cue (F_1,27_=1.31, p=.26, ƞ^2^_*p*_=0.05, BF_01_=2.95) and we also did not observe a significant 3-way interaction (F_2,54_=0.80, p=.46, ƞ^2^_*p*_=0.03, BF_01_=4.46; **Figure 4 – Supplement 1B**).

Third, and finally, similar to the effect of stimulus identity, drug condition affected ERP amplitudes relatively late in time (406-586ms; p=.004; **Figure 4C – right panel**). Specifically, ATX showed a trend toward lower ERP amplitudes compared to PLC (F_1,27_=3.56, p=.07, ƞ^2^_*p*_=0.12; **Figure 4D >– right panel**), whereas DNP increased amplitudes over occipito-temporal regions in this time-window (F_1,27_=4.86, p=.04, ƞ^2^_*p*_=0.15). We did not observe any significant interaction effects with drug condition and cue on ERP amplitude in this time-window (F_2,54_=0.37, p=.69, ƞ^2^_*p*_=0.01, BF_01_=6.43).

### Neuromodulatory effects on preparatory attention

The predictive cues in our task allowed participants to anticipate targets appearing at certain locations, by covertly shifting their locus of attention. Previous work has identified a neural marker for attentional orienting in response to a visual cue, i.e. an early negative amplitude deflection over contralateral regions (early directing attention negativity, EDAN; Murray et al., 2011; Praamstra & Kourtis, 2010). Moreover, attentional shifts have been associated with changes in cortical excitability, indexed by lateralized alpha-band (8-12Hz) power suppression (stronger alpha suppression at the contralateral side of the stimulus; Capotosto et al., 2009; Händel et al., 2011; Jensen & Mazaheri, 2010; Thut, 2006). Recent studies also suggests that neuromodulatory systems may be involved in regulating changes in global cortical excitability in relation to shifts in spatial attention (Bauer et al., 2012; Dahl et al., 2022). Here, we tested whether enhanced catecholaminergic and cholinergic levels modulated these electrophysiological markers of preparatory spatial attention. For the following EEG analyses, we used two symmetrical regions of interest (ROIs) over occipital areas that are commonly used for analyses regarding preparatory attention (left ROI: O1, PO3 and PO7; right ROI: O2, PO4 and PO8; Capotosto et al., 2009; Kelly et al., 2006; Thut, 2006).

In the next set of analyses, we will inspect prestimulus activity for the three different drug conditions, locked to the attentional cue. Therefore, there is no factor stimulus identity, but the logic defining the cue conditions is the same as illustrated in **Figure 4A**. The factor cue again had two levels and referred to the hemisphere associated with the cued location (the hemisphere contralateral to the cued direction) and the hemisphere associated with the uncued location (the hemisphere ipsilateral to the cued direction). For simplicity, we will refer to these factor levels as the cued (contra) and the uncued (ipsi) location from here on. We plotted the cue-locked ERPs for all 6 conditions in **Figure 5A** (cue x drug). Neural activity for the hemisphere associated with the cued location (contra) was clearly different than for the uncued location (ipsi) in two time-windows (89-175ms, p=.002; 198-363ms, p<.001). This analysis thus revealed a strong lateralization of neural activity due to the shift of attention towards the cued location. Interestingly, the early lateralized component, resembling the early directing attention negativity (EDAN), was modulated by drug (interaction: drug x cue, F_2,54_=5.05, p=.01, ƞ^2^_*p*_=0.16; **Figure 5B**) and this interaction was driven by ATX (ATX vs. PLC: F_1,27_=23.95, p<.001, ƞ^2^_*p*_=0.47; DNP vs. PLC: F_1,27_=1.47, p=.24, ƞ^2^_*p*_=0.05, BF_01_=2.12). Cue related lateralized activity in the later cluster showed a trending effect of overall drug modulation (F_2,54_=3.12, p=.05, ƞ^2^_*p*_=0.10; **Figure 5 – Supplement 1A**), but pairwise post-hoc tests showed no reliable differences for ATX/DNP vs. PLC when tested separately (ATX: F_1,27_=1.44, p=.24, ƞ^2^_*p*_=0.05, BF_01_=2.31; DNP: F_1,27_=1.32, p=.26, ƞ^2^_*p*_=0.05, BF_01_=2.61).

**Figure 5.**
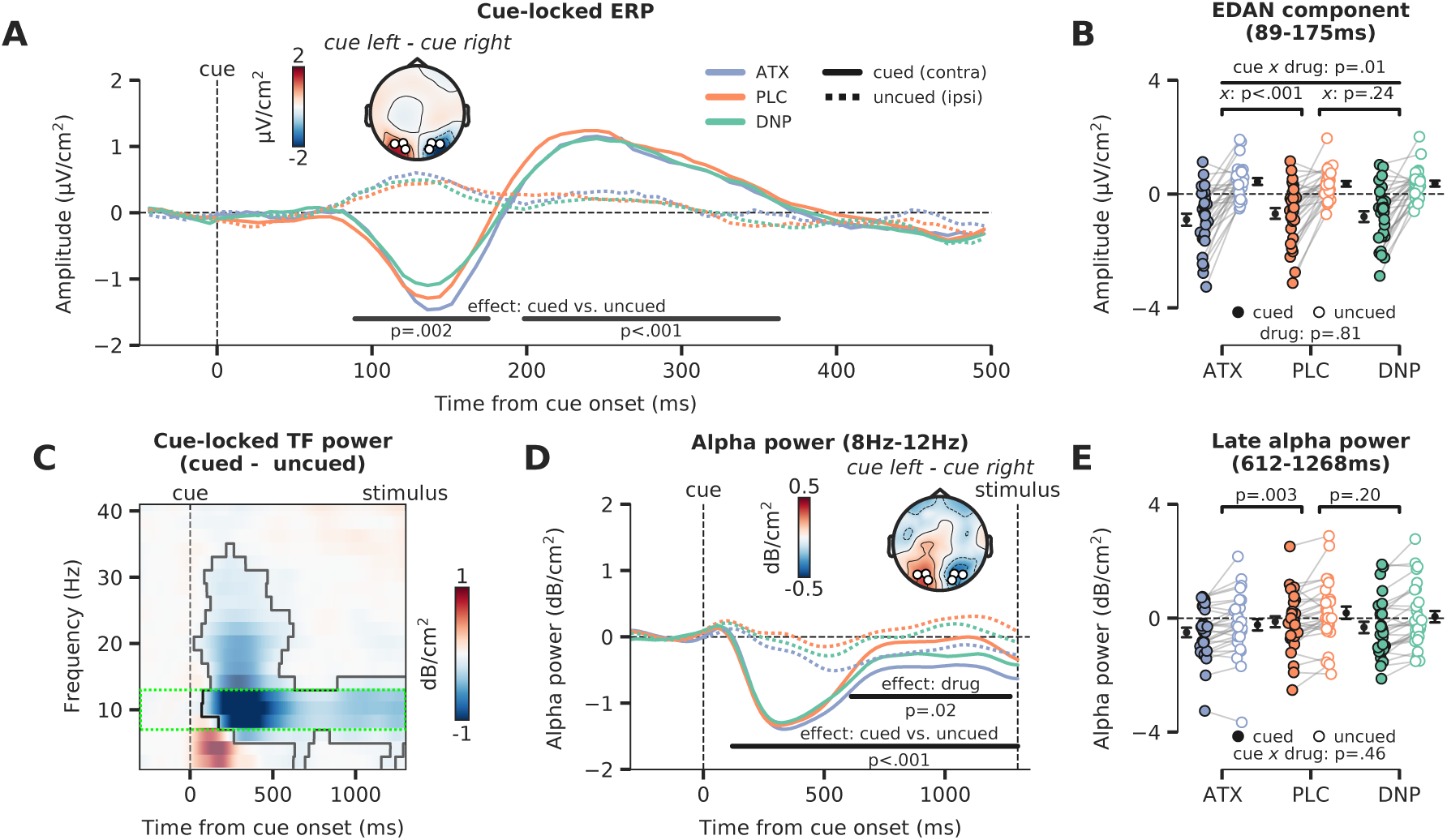
Attentional and catecholaminergic modulation of prestimulus cortical activity. **A)** Cue-locked ERP (µV/cm^2^) over occipital regions reflecting the cued location (hemisphere contralateral to the cued direction) and reflecting the uncued location (hemisphere ipsilateral to the cued direction), split up for the three drug conditions. Black horizontal bars show rmANOVA main effects of cue, cued (contra) vs. uncued (ipsi), cluster corrected for multiple comparisons. The topographic map shows the contrast between a cue to the left vs. a cue to the right (time-window: 89-175ms, early cluster in which we observed the lateralization effect). White markers indicate the ROIs used for all cue-locked analyses. **B)** Average activity within the early temporal cluster for the hemisphere associated with the cued (contra) vs uncued (ipsi) location as a factor of drug, showing lateralized effects. The p-value of the interaction effect between cue and drug (pairwise vs. PLC) is demarked with *x*. **C)** The difference in time-frequency (TF) power (dB/cm^2^) for cued (contra) vs uncued (ipsi) locations (lateralization effect). The significant TF cluster (highlighted by the solid black line) is derived from cluster-based permutation testing controlling for multiple comparisons. The green dotted box indicates the alpha-band time-frequency ROI that is used to compute panel **D**. **D)** Cue-locked prestimulus alpha power (dB/cm^2^) over occipital regions associated with the cued (contra) and uncued (ipsi) location, split up for the three drug conditions. Black horizontal bars show rmANOVA main effects for cue (contra vs. ipsi, lateralization effect) and drug. The topographic map shows the contrast between when a cue to the left vs. a cue to the right was presented (time-window: 659-1253ms, cluster in which we observed a main effect of drug). **E)** Average alpha power within the late temporal cluster in which drug effects on alpha power were observed, split up for cue and drug conditions. The p-values on top reflect the main effects between ATX/DNP and PLC and the p-value at the bottom demarked with *x* depicts the omnibus interaction effect between drug condition and cue.

Next, we focused on alpha-band dynamics from the onset of the attentional cue leading up to stimulus presentation (1300ms post-cue). Replicating previous work, the time-frequency (TF) spectrum plotted in **Figure 5C** shows that attentional cues resulted in a stronger suppression of alpha-band power for the hemisphere associated with the cued location (contra) compared to the uncued (ipsi) location. To relate these effects to drug condition, we plotted alpha power (8-12 Hz) for the hemisphere associated with the cued and uncued locations for all 3 drug conditions separately (**Figure 5D)**. Stronger alpha-band desynchronization for cued vs uncued locations was observed 120ms from cue onset until target stimulus onset (p<.001; **Figure 5D**). Moreover, there was a main effect of drug, revealing that total alpha power over both the hemisphere associated with the cued location and the hemisphere associated with the uncued location, was modulated by drug from 659-1253ms after cue onset (p=.02; **Figure 5D**). Again, this main effect of drug was driven by catecholaminergic activity (ATX: F_1,27_=10.42, p=.003, ƞ^2^_*p*_=0.28; DNP: F_1,27_=1.73, p=.20, ƞ^2^_*p*_=0.06, BF_01_=1.30; **Figure 5E**). Drug condition did not modulate the effect of cue on prestimulus-alpha power (F_2_,=10.42, p=.46, ƞ^2^_*p*_=0.03, BF_01_=4.99; **Figure 5E**). This latter result shows that ATX reduced overall alpha-band power for both cued (contra) and uncued (ipsi) locations, with respect to PLC, so not in a lateralized manner. The effect of the spatial cue on prestimulus alpha-band power lateralization in this late time window was related to the cue effect on perceptual sensitivity and this relation was not different between the drug sessions (see **Figure 5 - Supplement 1B** for details).

In sum, replicating a wealth of previous attention studies, both occipital ERPs (EDAN) as well as occipital alpha-band suppression were affected by shifting the locus of spatial attention, as reflected in clear neural differences between activity measured over the hemisphere associated with cued (contra) vs. the uncued (ipsi) location. Interestingly, ATX increased the lateralized cue-locked ERP component (EDAN) by enhancing the difference between cued (contra) and uncued (ipsi) locations, suggesting enhanced attentional orientation in a hemisphere specific manner **(Figure 5A**). However, in contrast, prestimulus occipital alpha power, an index of cortical excitability (Capotosto et al., 2009; Händel et al., 2011; Jensen & Mazaheri, 2010), was bilaterally decreased under ATX, and did not relate to the attentional cue, suggesting instead that enhanced catecholaminergic levels regulate cortical excitability in a spatially non-specific and global manner (**Figure 5C**).

## Discussion

In this study, we used a 3x2 factorial design in which we manipulated neuromodulator levels (through pharmacology) and spatial attention, allowing us to causally demonstrate their separate, shared, and interactive effects on perceptual decision-making in healthy human participants. By administering atomoxetine (ATX) and donepezil (DNP), we were further able to tease apart the roles of cholinergic and catecholaminergic neuromodulatory systems respectively, in relation to, and in interaction with, spatial attention.

### Spatial attention and catecholaminergic neuromodulation jointly shape perceptual decision-making

Computational modelling of choice behavior showed that both the allocation of attention to target stimuli (rotated Gabor patches) and increased catecholaminergic levels enhanced the rate of sensory evidence accumulation (drift rate, **Figure 2D**), resulting in improved perceptual sensitivity (effect was stronger for attention, **Figure 2A)**. In other words, participants were better able to tell whether a presented Gabor stimulus was oriented clockwise or counterclockwise when Gabors were attended (validly cued vs. invalidly cued) and when atomoxetine was received (vs. placebo). These effects were reflected in a well-known neural marker of sensory evidence accumulation measured over parietal cortex (CPP, **Figure 3D**), as elevated catecholaminergic levels increased CPP peak amplitude and attention increased both its slope and its peak amplitude. Although the effects of attention and catecholaminergic neuromodulation on algorithmic and neural markers of sensory evidence accumulation were ostensibly similar, additional EEG analyses revealed that attention and catecholamines differently modulated activity over occipito-temporal regions. First, we observed that the effects of attention and catecholaminergic neuromodulation on stimulus-locked (Gabor) neural activity over occipito-temporal electrodes occurred at different latencies. Attention modulated sensory processing much earlier in time (starting ∼70ms after stimulus presentation) than the catecholaminergic intervention (starting ∼400 after stimulus presentation; **Figure 4A**). Second, both spatial attention and catecholamines modulated alpha-band desynchronization, a marker of cortical excitability (**Figure 5D**), but attention did so unilaterally, as reflected in stronger alpha-band desynchronization in the hemisphere opposite to the locus of attention, whereas catecholaminergic modulation of alpha-band desynchronization was bilateral (no hemispherical specificity nor interaction with attentional cue). Both findings are in line with recent work showing independent modulations of spatial attention and arousal on neuronal activity in mouse V1 (Kanamori & Mrsic-Flogel, 2022). Thus, prestimulus cortical excitability (alpha-power) and stimulus-locked activity related to the processing of visual input were differentially modulated by spatial attention and catecholaminergic neuromodulation.

We did observe two striking interaction effects between the catecholaminergic system and spatial attention. First, effects of attention on drift rate were increased under catecholaminergic enhancement (**Figure 2D**). Although this interaction effect was not reflected in CPP slope/peak amplitude, this does suggest that catecholamines and spatial attention might together shape sensory evidence accumulation in a non-linear manner. Second, the amplitude of the cue-locked early lateralized ERP component (resembling the EDAN) was increased under ATX as compared to PLC. The underlying neural processes driving the EDAN ERP, as well as its associated functions, have been a topic of debate. Some have argued that the EDAN reflects early attentional orienting (Praamstra & Kourtis, 2010) but others have claimed it is mere a visually evoked response and reflects visual processing of the cue (Velzen & Eimer, 2003). Thus, whether this effect reflects a modulation of ATX on early attentional processes or rather a modulation of early visual responses to sensory input in general is a matter for future experimentation.

Taken together, spatial attention and neuromodulation showed similar, unique and interactive effects on neural activity underlying different stages of the perceptual decision-making process. This intriguing relation between spatial attention and neuromodulation advances our understanding about how these systems jointly shape perceptual decision-making but also indicates the necessity for further exploration of the complex interactions between these systems.

### Tonic versus phasic neuromodulation in relation to attentional processes

In contrast with recent work associating catecholaminergic and cholinergic activity with attention by virtue of modulating prestimulus alpha-power shifts (Bauer et al., 2012; Dahl et al., 2020, 2022) and attentional cue-locked gamma-power (Bauer et al., 2012; Howe et al., 2017), the current work shows that the effects of neuromodulator activity are relatively global and non-specific, whereas the effects of spatial attention are more specific to certain locations in space. Our findings are, however, not necessarily at odds with these previous studies. Most recent work associates phasic (event-related) arousal with selective attention (for reviews see: Dahl et al., 2022; Thiele & Bellgrove, 2018). For example, cue detection in visual tasks is known to be related to cholinergic transients occurring after cue onset (Howe et al., 2017; Parikh et al., 2007). Contrarily, in our work we aimed to investigate the effects of increased baseline levels of neuromodulation by suppressing the reuptake of catecholamines and the breakdown of acetylcholine throughout cortex and subcortical structures. Tonic and phasic neuromodulation have previously been shown to differentially modulate behavior and neural activity (de Gee et al., 2014, 2020, 2021; McGinley et al., 2015; McGinley, Vinck, et al., 2015; van Kempen et al., 2019). Note, however, that it is difficult to investigate causal effects of tonic neuromodulation in isolation of changes in phasic neuromodulation, mostly because phasic and tonic activity are thought to be anti-correlated, with lower phasic responses following high baseline activity and vice versa (Aston-Jones & Cohen, 2005; de Gee et al., 2020; Knapen et al., 2016). As such, pharmacologically elevating tonic neuromodulator levels may have resulted in changes in phasic neuromodulatory responses as well. Concurrent and systematic modulations of tonic (e.g. with pharmacology) and phasic (e.g. with accessory stimuli; Bruel et al., 2022; Tona et al., 2016) neuromodulator activity may be necessary to disentangle the respective and interactive effects of tonic and phasic neuromodulator activity on human perceptual decision-making.

### Relating the algorithmic and neural implementations of evidence accumulation

We reported attentional and neuromodulatory effects on algorithmic (DDM, **Figure 2**) and neural (CPP, **Figure 3**) markers of sensory evidence accumulation. Recent work has started to investigate the association of these two descriptors of the accumulation process, aiming to uncover whether neural activity over centroparietal regions reflects evidence accumulation, as proposed by computational accumulation-to-threshold models (Kelly & O’Connell, 2015; O’Connell et al., 2018; O’Connell & Kelly, 2021; Twomey et al., 2015). Currently, the CPP is often thought to reflect the decision variable, i.e. the (unsigned) evidence for a decision (Twomey et al., 2015), and consequently its slope should correspond with drift rate, whereas its amplitude at any time should correspond with the so-far accumulated evidence. As -computationally-the decision is reached when evidence crosses a decision bound (the threshold), it may be argued that the peak amplitude of the CPP (roughly) corresponds with the decision boundary. This seems to contradict our observation that 1) ATX modulated drift rate, but not CPP slope and 2) ATX did not modulate boundary separation, but did modulate CPP peak. Note, however, that previous studies using pharmacology or pupil-linked indexes of (catecholaminergic) neuromodulation have also demonstrated effects on both CPP peak (van Kempen et al., 2019) and CPP slope (Loughnane et al., 2019).

Here, we demonstrated that response accuracy and response speed are differentially represented in the CPP, with correct vs. erroneous responses resulting in a higher slope and peak amplitude, whereas fast vs. slow responses are only associated with increased slopes (**Figure 3A-B**). Speculatively, the specific effect of any (pharmacological) manipulation on the CPP may depend on task-setting. For example, Loughnane et al. (2019) used a visual task on which participants did not make many errors (hit rate>98%, no false alarms), whereas we applied a task in which participants regularly made errors (roughly 25% of all trials). Possibly, the effects of ATX from Loughnane et al. (2019) in terms of behavior (RT effect, not accuracy/d’) and CPP feature (slope effect, not peak) may therefore have been different from the effects of ATX we observed on behavior (d’ effect, not RT) and CPP feature (peak effect, not slope). Regardless, when we compared subjects with high and low drift rates (**Figure 3C**), we observed that both CPP slope and CPP peak were increased for the high vs. low drift group (independent of the drug or attentional manipulation). This indicates that both CPP slope and CPP peak were associated with drift rate from the DDM. Clearly, more work is needed to fully understand how evidence accumulation unfolds in neural systems, which could consequently inform future behavioral models of evidence accumulation as well.

### Attentional and neuromodulatory effects on sensory processing or decision-making?

Fluctuations in arousal states have been both associated with alterations in sensory representations (Podvalny et al., 2021; Vinck et al., 2015; Warren et al., 2016) and modulations of response criterion or other latent variables of the decision process (de Gee et al., 2014, 2017, 2020). Similarly, human work on alpha-band dynamics −often used as a proxy for attentional state− has not yet been able to arbitrate between the hypotheses that decreased (unlateralized) alpha-band power may reflect an overall increase in baseline excitability (resulting in amplified responses to both signal and noise) or whether baseline excitability may decrease/increase the trial-by-trial precision of neural responses (Samaha et al., 2020). The first case would result in alterations in response criterion (i.e. decision-making), whereas the latter case would lead to less/more accurate perceptual performance and sensitivity (i.e. likely related to perception) (Busch et al., 2009; Iemi & Busch, 2018; Samaha et al., 2017, 2020; VanRullen et al., 2011; Zhou et al., 2021). Here we provide evidence that cholinergic and catecholaminergic neuromodulation seem to affect stimulus-locked neural responses relatively late in time (close to the execution of the motor response), whereas the effects of attention were observable much earlier. This corroborates earlier findings showing, respectively, late and early effects of neuromodulation and attention on cortical activity (Alilović et al., 2019; Baumgartner et al., 2018; Gelbard-Sagiv et al., 2018; Hillyard & Anllo-Vento, 1998; Loughnane et al., 2019; Murphy et al., 2011; Nuiten et al., 2021; Poort et al., 2012). Although, based on the timing of these effects, it might be tempting to conclude that attention primarily modulates (early) perceptual processes and neuromodulation (late) decisional processes, this conclusion is at present premature. Our study was not set-up to dissociate perceptual from decisional effects, as these are hard to disentangle in practice (see also Sánchez-Fuenzalida et al., 2022), and cannot be isolated based on timing alone (Alilović et al., 2023). However, this study highlights that various cortical states – e.g. attention and arousal– may distinctly affect stimulus-locked cortical activity and therefore need to be considered jointly when addressing the effect of (prestimulus) cortical state variations on perceptual decision-making (Podvalny et al., 2021; Waschke et al., 2019).

### Elevated cholinergic levels modulate neural activity in absence of bodily and behavioral effects

In line with previous non-clinical studies that report physiological responses of 5mg DNP (Pfeffer et al., 2018, 2021), we did not observe consistent subjective or physiological effects of DNP. However, even in the absence of such autonomic markers, 5mg DNP has been shown to induce behavior and neural activity, related to visual perception (Boucart et al., 2015; Gratton et al., 2017; Pfeffer et al., 2018, 2021; Silver et al., 2008). Although DNP did not significantly affect perceptual sensitivity and/or drift rate, we did observe a set of consistent neural effects, generally reflected in neural responses in the opposite direction as for ATX. This was the case for the CPP peak and slope (**Figure 3D-F**) as well as for occipito-temporal signals related to stimulus and cue processing (**Figure 4A/C**, **Figure 5A-B**). Corresponding with our observation that DNP and ATX effects were mirrored, a recent study using DNP and ATX in combination with whole-brain fMRI revealed that these neuromodulator systems have opposite effects on cortex-wide connectivity patterns, even though both systems enhanced neural gain (Pfeffer et al., 2021). More specifically, catecholamines enhanced and acetylcholine suppressed cortex-wide auto-correlation of activity (Pfeffer et al., 2021), suggested to reflect intracortical signaling by lateral and feedback processes (Hasselmo & Sarter, 2011; Silver et al., 2008). Furthermore, high cholinergic levels are associated with enhanced afferent input to cortex (feedforward) but decreased feedback drive (Hasselmo & Sarter, 2011). Because both evidence accumulation processes and target selection processes strongly rely on cortical feedback (Boehler et al., 2009; Dehaene et al., 2011; Donohue et al., 2020; Lamme & Roelfsema, 2000; Murphy et al., 2021; Pereira et al., 2022; Pitts et al., 2014; Seijdel et al., 2021; Theeuwes, 2010; Wang, 2008), the mirrored (compared to ATX) effects of DNP on the neural markers of these processes may have been caused by interrupted feedback processing. Furthermore, although previous work has implicated the cholinergic system in prestimulus alterations in neural excitability (Bauer et al., 2012), indexed by alpha-power desynchronization, we did not find evidence for this (**Figure 5E**), possibly because Bauer et al. (2012) used physostigmine, a more potent cholinergic esterase inhibitor than DNP (Ogura et al., 2000). Future work applying various stimulants and suppressors of cholinergic activity, administered in different dosages, is necessary to firmly establish the role of the cholinergic system in controlling cortical functioning and spatial attention.

### Limitations of the current study

Although the effects of the attentional manipulation were generally strong and robust, the statistical reliability of the effects of the pharmacological manipulation was more modest for some comparisons. This may partly be explained by high inter-individual variability in responses to pharmaceutical agents. For example, initial levels of catecholamines may modulate the effect of catecholaminergic stimulants on task performance, as task performance is supposed to be optimal at intermediate levels of catecholaminergic neuromodulation (Cools & D’Esposito, 2011). While acknowledging this, we would like to highlight that many of the observed effects of ATX were in the expected direction and in line with previous work. First, pharmacologically enhancing catecholaminergic levels have previously been shown to increase perceptual sensitivity (d’) (Gelbard-Sagiv et al., 2018), a finding that we have replicated here. Second, methylphenidate (MPH), a pharmaceutical agent that elevates catecholaminergic levels as well, has been shown to increase drift rate as derived from drift diffusion modeling on visual tasks (Beste et al., 2018) in line with our ATX observations. Third, a previous study using ATX to elevate catecholaminergic levels observed that ATX increased CPP slope (Loughnane et al., 2019). Although in our case ATX increased the CPP peak and not its slope, this provide causal evidence that centro-parietal ERP signals related to sensory evidence accumulation are modulated by the catecholaminergic system (Nieuwenhuis et al., 2005). Fourth, we observed that elevated levels of catecholamines affected stimulus driven occipital activity relatively late in time and close to the behavioral response, which resonates with previous observations (Gelbard-Sagiv et al., 2018). Finally, ATX had robust effects on physiological responses (heart rate, blood pressure, pupil size), cue-locked ERP signals and oscillatory power dynamics in the alpha-band, leading up to stimulus presentation. We concur, however, that more work is needed to firmly establish how (various forms of) attention and catecholaminergic neuromodulation affect perceptual decision-making.

## Conclusion

Summarizing, we show that elevated catecholaminergic levels and spatial attention jointly shape perceptual decision-making. Similar, unique, and interactive effects of attention and catecholaminergic neuromodulation on behavioral, algorithmic, and neural markers of decision-making were observed. Together, this study reveals an intricate, complex, and yet to be further explored, relationship between the attentional and catecholaminergic systems.

## Author contributions

Conceptualization: SAN, SVG

Medical assessment of participants: JBZ

Data collection: SAN

Data analysis: SAN, JWG

Writing – original draft: SAN, SVG

Writing – review & editing: JWG, JJF

## Data and materials availability

All data and analysis scripts will be made available on FigShare upon acceptance.

## STAR Methods

### Participants

For this experiment 30 male, Dutch-speaking, human participants were recruited from the online research environment of the University of Amsterdam. All participants were aged between 18 and 30. Given the pharmacological nature of this experiment, participants were only included after passing extensive physical and mental screening (see **Screening procedure**). This study was approved by the Medical Ethical Committee of the Amsterdam Medical Centre (AMC) and the local ethics committee of the University of Amsterdam. Written informed consent was obtained from all participants after explanation of the experimental protocol. Two participants decided to withdraw from the experiment after having performed the first experimental session. The data from these participants are not included in this work, resulting in N=28. Participants received monetary compensation for participation in this experiment.

### Screening procedure

Following registration, candidate participants were contacted via e-mail to inform them of the inclusion criteria, exact procedures, and possible risks. In addition, candidate participants were provided with the credentials of an independent physician that was available for questions and further explanation. The participants were contacted after a deliberation period of seven days to invite them for a pre-intake via telephone. During this pre-intake, it was ensured that the candidate participant indeed met all inclusion and exclusion criteria. If so, participants were invited for an intake session at the research facility of the University of Amsterdam. During this in-person intake, the experimental protocol, including the following physical and mental assessment, was explained in detail after which candidate participants provided written consent. The intake further consisted of a set of physiological measures -body mass index (BMI), heart rate, blood pressure, and an electrocardiogram (ECG)- and a psychiatric questionnaire to assess mental health. The data from the intake were assessed by a physician, who subsequently decided whether a candidate participant was eligible for participation. Lastly, participants performed the staircasing procedure for the behavioral task (see **Staircasing procedure**).

### Drug administration

The study was conducted using a randomized, double-blind crossover design. Atomoxetine (40mg), donepezil (5mg) and placebo were administered in different experimental sessions, separated by at least seven days. The order of drug administration was counterbalanced between participants. Experimental days started at 09:00 and ended at 16:00. Atomoxetine (∼2 hours) and donepezil (∼4 hours) reach peak plasma levels at different latencies, therefore these pharmaceuticals were administered at different times prior to the onset of the behavioral tasks (**Figure 1A**). To ensure the double-blind design, participants were required to orally ingest a pill 4 hours prior to the onset of the first behavioral task, which could contain either donepezil or a placebo, and a second pill 2 hours prior to onset of the first behavioral task, which could contain either atomoxetine or a placebo. Thus, participants received one placebo and a working pharmaceutical or two placebos on every experimental session.

The pharmaceuticals used in this study were chosen based on their pharmacokinetic and pharmacodynamic properties, the relatively limited side-effects, and prior use of these pharmaceuticals in other studies in the cognitive sciences (Boucart et al., 2015; Pfeffer et al., 2018, 2021; Warren et al., 2016). Atomoxetine is a relatively selective noradrenaline reuptake inhibitor, which inhibits the presynaptic noradrenaline reuptake transporter, thereby resulting in increased noradrenaline and dopamine levels in the synaptic cleft (Simpson & Plosker, 2004). The half-life of atomoxetine varies between 4.5-19 hours and peak plasma levels are reached ∼2 hours after administration. Donepezil is a cholinesterase inhibitor, which impedes breaking down of acetylcholine by cholinesterase thereby resulting in increased acetylcholine levels in the synaptic cleft (Rogers & Friedhoff, 1998). The elimination half-life of donepezil is 70 hours and peak plasma levels are reached after ∼4 hours.

Minor side-effects were reported by our participants, including a mild headache (1 DNP session, 2 ATX session) and mild nausea (1 DNP session, 4 ATX sessions). Other side-effects, such as fatigue and tenseness, were equally present across all drug sessions including PLC.

### Design and procedures

#### Experimental setting

Participants performed several tasks during a single experimental day. The first task was an auditory discrimination/detection task, lasting two hours, that was executed directly after the administration of the first pill. The data of that experiment fall outside the scope of this paper. An EEG apparatus was connected 3.5 hours after ingestion of the first pill, ensuring that participants could start the main experiment precisely 4 hours after ingestion of the first pill, supposedly when blood concentration levels for both atomoxetine and donepezil were peaking. During this main part of the experiment participants performed five different computerized visual perception tasks, out of which we will only discuss the spatial attention task. The order of these behavioral tasks was counterbalanced between participants but was maintained over sessions. Participants were seated in a darkened, sound isolated room, resting their chins on a head-mount located 80cm from a 69x39cm screen (frequency: 60Hz, resolution: 1920 x 1080). The main task and staircase procedure were programmed in Python 2.7 using PsychoPy (Peirce, 2007) and in-house scripts.

#### Cued orientation discrimination paradigm

Participants performed a task that was adapted from the Posner cueing paradigm (Posner, 1980). For the task, participants were asked to report the orientation of Gabor patches as being clockwise (45°) or counterclockwise (-45°), by pressing left (“s” on a keyboard, for counterclockwise) or right (“k”, for clockwise) with their left or right index finger respectively. We did not specifically instruct participants to respond as quickly or accurately as possible. The Gabor patches were presented for 200ms on either the left or right side of the screen (center at 8°, radius 5.5°, spatial frequency 1.365 cycles/degree). Simultaneously, circular patches containing dynamic noise were presented bilaterally (center at 8°, radius 6.5°). Prior to stimulus presentation, a visual cue was presented for 300ms near the fixation mark (horizontal dash, center at 0.33°). This cue was predictive of the stimulus location with a cue validity of 80%, meaning that the cue matched the stimulus location in 80% of the trials. Participants were instructed to use the cue, by covertly shifting attention (i.e. without moving eyes) towards the cued location. In 20% of trials the cue was invalid and the stimulus would appear on the opposite side. After cue offset there was a 1000ms interval until stimulus onset. The response window started concurrently with stimulus presentation and lasted 1400ms. If participants did not respond during this window they would receive visual feedback informing them of their slow response (in Dutch: “te laat”, which translates into: “too late”). A variable inter-trial interval (ITI) of (250ms-350ms) started directly after a response or the end of the response-window. Stimulus locations, stimulus orientation and cue direction were all balanced to occur on 50% of the trials (i.e. 50% counterclockwise, 50% left location, 50% cue to the right etc.). The task was divided in two blocks of 280 trials, which in turn were subdivided in shorter blocks of 70 trials, in between which the participants could rest.

To ensure participants did not shift their gaze towards either the left or right, we measured gaze position via eyetracking (see **eyetracking**). Trials only commenced when participants’ gaze was at fixation (cutoff: 1.5°) and whenever participants lost fixation or blinked during a trial, the fixation mark would turn white and the data for the trial would be marked as faulty. At the end of each block, a new trial was presented for every trial that was terminated because of blinks or lost fixation. In total, participants performed 560 trials of this task without losing fixation or blinking. After 280 trials on which fixation was not lost, the eyetracker was recalibrated (see **Eyetracking)**.

#### Staircasing procedure

Participants performed a staircasing procedure during their intake session. The staircase task was almost identical to the primary task, but without predictive cues prior to stimulus onset. The stimulus properties, presentation time and response window were the same. The ITI was prolonged to 1100ms-1300ms. Participants received feedback on their performance on a trial-by-trial basis; the fixation dot turned green for correct answers and red for incorrect answers. An adaptation of the weighted up-down method proposed was used to staircase performance at 75% correct, by changing the opacity of the Gabor patch (Kaernbach, 1991). In short, opacity adjustment after erroneous responses were weighted differently than after correct responses, in a ratio of 3:1. The step size was 0.01 (opacity scale: 0-1, 1 is fully opaque), thus after errors the opacity would be increased by 0.01 and after correct answers it would be decreased by 0.01/3. The procedure was aborted after 50 behavioral reversals, i.e. changes in sequences from correct to error or vice versa. The output difficulty of the staircase procedure was calculated as the average opacity on reversal trials. In total, participants performed two blocks of this staircase procedure. The first block started at a high opacity, allowing the participants to become familiar with the stimulus. The second block started at the opacity obtained from the first block. After completing the staircasing procedure, task difficulty was fixed to be able to compare the effects of the pharmacological manipulation across all experimental sessions.

### Data acquisition and preprocessing

#### Eyetracking

Gaze position and pupil size were measured with an EyeLink 1000 (SR Research, Canada) eyetracker during the experiment at 1000Hz. Gaze position was moreover monitored online to ensure participants’ gaze remained at or near fixation (within 1.5°, horizontal axis). A nine-point calibration was used at the start of each block. To minimize movement of the participant, we used a head-mount with chinrest. Throughout the experiment participants were instructed to move their heads as little as possible and to try to refrain from blinking during a trial. Pupil traces were bandpass filtered between 0.01Hz-10Hz, blinks were linearly interpolated and the effects of blinks and saccades on pupil diameter were removed via finite impulse-response deconvolution (Knapen et al., 2016).

#### EEG acquisition, preprocessing and time-frequency decomposition

EEG-data were recorded with a 64-channel BioSemi apparatus (BioSemi B.V., Amsterdam, The Netherlands), at 512Hz. Vertical eye-movements were recorded with electrodes located above and below the left eye, horizontal eye-movements were recorded with electrodes located at the outer canthi of the left and the right eye. All EEG traces were re-referenced to the average of two electrodes located on the left and right earlobes. The data were high-pass filtered offline, with a cut-off frequency of 0.01Hz. Next, bad channels were detected automatically via a random sample consensus algorithm (RANSAC), implemented in the Autoreject Python package (Jas et al., 2017), and subsequently interpolated via spline interpolation. Next, epochs were created by taking data from -2000ms to 2000ms around onset of stimulus presentation. To remove eyeblink artefacts, an independent component analysis (ICA; 25 components) was performed on the epoched data and components that strongly correlated to vertical EOG data were excluded. On average 1.23 (sd: 0.47, maximum: 3) components were rejected per file. Remaining artefacts were automatically detected by using the same RANSAC algorithm as before but on epoched data. Bad segments were repaired via interpolation if the artifactual data was present in only a few channels, but if more channels were affected the epoch was removed from the EEG data. On average 3.82% (sd: 5.16, maximum: 24.18%) of all epochs were removed. Lastly, the scalp current density (CSD) was computed using the surface Laplacian to attenuate the effects of volume conductance (Cohen, 2014).

Time-frequency (TF) representations of EEG data were calculated from epochs that were extracted from a time-window -1600ms to 0ms prestimulus. Time-frequency power was obtained through convolution of Morlet wavelets with epoched data from 2-40Hz in steps of 2Hz. For each frequency, we defined the number of cycles as:

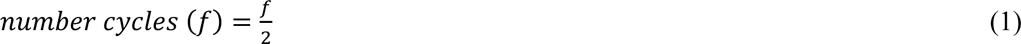

Then we averaged TF data across cue validity conditions and spatial locations (four stimulus categories). Next, both this average TF power, as well as single trial TF power, was normalized to a baseline of -250ms to -50ms pre-cue with a decibel (dB) transformation:

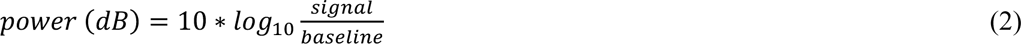

### Data analysis

All behavioral analyses were programmed in Python 3.7. Trials during which fixation was lost missed trials and trials with a reaction time (RT) larger than 1400ms were excluded from all analyses. The remaining data was used for the main behavioral analyses, but for the alpha power binning analysis of **Figure 5 – Supplement 2,** only behavioral data belonging to unrejected EEG epochs were included to match single trial EEG to behavior. EEG analyses were performed with use of the Python package MNE (version 0.24.0; Gramfort et al., 2013)

#### Statistical analysis of physiological data, side-effects, and self-reports

We obtained bodily (heart rate and blood pressure) and subjective (Visual Analogue Scale, VAS; Bond & Lader, 1974) measurements of arousal on three occasions during each session; before drug intake (baseline), prior to the onset of the first EEG task (after four hours) and at the end of each session (after seven hours). We calculated mean arterial pressure (MAP) from systolic and diastolic blood pressure as:

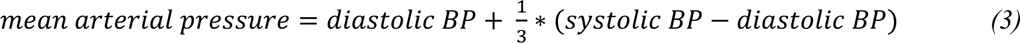

VAS scores were calculated by measuring the location of marked answers in centimeters from the left. Next, all 16 scores of the VAS were into three categories: contentedness, calmness and alertness following previous methods (Bond & Lader, 1974). HR, MAP and VAS scores were corrected by calculating percentage change from the baseline measurement. Next, we applied 1-factor rmANOVAs (factor drug), on these scores for each of the remaining timepoints. Post-hoc, pairwise t-tests were performed to test the effects of ATX and DNP vs. PLC.

We assessed the effects of drug on pupil diameter right before the onset of the behavioral tasks (at t=4 hours). We defined minimal (bright monitor background) and maximal (dark monitor background) pupil diameter as the average pupil size within the final 5s of each presentation window of 15s. Next, we performed similar rmANOVAs and post-hoc t-tests on these raw (non-normalized) pupil diameters. To test the occurrence of side effects as well proportion of forced guesses of drug intake at the end of each session, we used binomial tests. Specifically, we calculated proportions of side-effects and guesses about the nature of drug intake (placebo or pharmaceutical) under PLC. Then we calculated the same proportions for ATX and DNP and tested them against the PLC proportion with a binomial test.

#### Statistical analysis of behavioral data

To investigate behavioral performance on this task, we used perceptual sensitivity (d’) and bias (criterion) derived from signal detection theory (SDT; Green & Swets, 1966) in addition to RTs. To test the effect of cue validity and drugs on these measures we performed 3x2 factorial rmANOVAs (factors: drug and cue validity). We did not include drug order as a between-subject variable in our statistical models, because drug order was counter-balanced between participants. Bayesian rmANOVAs (uniform prior, default setting) were used in the case of null-findings, to test support for the null-hypothesis (JASP, 2022). We expressed all effect sizes as ƞ^2^_*p*_. For one-tailed t-tests we calculated these as follows:

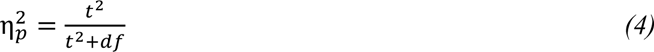

Where *t* is the t-statistic and df the degrees of freedom. Effect sizes are considered small when ƞ^2^_*p*_=0.01, medium when ƞ^2^_*p*_=0.06 and large when ƞ^2^_*p*_=0.14 (Cohen, 1988).

#### Drift diffusion modelling

We constructed a drift diffusion model (DDM) to gain insight into which parameters of the decision process were modulated by attention and drug condition (Ratcliff & McKoon, 2008). Specifically, we used the Python package HDDM to fit hierarchical Bayesian accuracy-coded regression models to reaction time distributions of correct and incorrect trials for every participant (Wiecki et al., 2013). In a first regression model, we allowed drift rate, non-decision time and decision bound separation to vary with drug, cue validity and their interaction. To estimate the effects of drug, we fitted these regression models separately for ATX (vs. placebo) and DNP (vs. placebo). We applied a weighted effect coding scheme, meaning that we coded our regressors as -1 and 1 (instead of dummy coding: 0 and 1). We weighted the regressors for cue validity according to the proportion of valid and invalid trials (-0.8 for invalid, 0.2 for valid), because valid and invalid trial counts were unbalanced. The effects of this model can be interpreted similarly as effects derived from an ANOVA. The Bayesian implementation of this model constrained single subject parameter estimates based on population estimates, making the model more robust to noise. Note that we also fitted unweighted effect coded versions of these models to verify the output of our weighted models, as well as two additional (weighted effect-coded) models to test whether drift rate variability was also modulated by cue validity and drug condition (**Figure 2 – Supplements 1-2**; Murphy et al., 2014).

#### EEG - ERP analysis

All analyses were performed on the CSD-transformed EEG data. To assert how spatial attention and elevated baseline levels of neuromodulators shape visual perception, we looked at neural activity related to the processing of the visual input (cue and stimulus) as well as activity related to the sensory information accumulation process preceding the response. For statistical analyses, epoched data were downsampled to 128Hz. We used distinct spatial ROIs for each of these events, based on previous literature. For cue processing, we used symmetrical electrode pairs O1/O2, PO3/PO4, and PO7/PO8 (Capotosto et al., 2009; Kelly et al., 2006; Thut, 2006), for processing of the visual stimulus, we used the symmetrical electrode pairs P7/P8 and P9/P10 (Loughnane et al., 2016; Newman et al., 2017; Papaioannou & Luck, 2020; van Kempen et al., 2019) and for response-locked evidence accumulation signals (i.e. centroparietal positivity, CPP) we used electrodes CPz, Cp1, and Cp2 (O’Connell et al., 2012; van Kempen et al., 2019; van Vugt et al., 2019). To extract ERPs, we first normalized epochs by subtracting the average baseline activity -80ms to 0ms before cue onset (for cue-locked ERP) and stimulus onset (for stimulus-locked and response-locked ERP).

We calculated the peak amplitude of the CPP as the amplitude at the time of the response and the slope of the CPP by fitting a linear regression in a time-window ranging from -250ms to 0ms before the response. Note that the time-window used for CPP slope estimation was chosen on the basis of visual inspection of the grand average CPP and roughly coincided with previous time-windows used for slope estimation (van Kempen et al., 2019; van Vugt et al., 2019). Next, CPP peak amplitude and slope were tested for incorrect vs. correct and fast vs. slow response trials (median split) with two-sided pairwise t-tests and for high vs. low drift rate participants with independent t-tests (**Figure 3A-C**). Moreover, 2x3 (attention x drug) rmANOVAs and post-hoc two-sided t-tests (drug vs. placebo) were used to test the effects of cue validity and drug on CPP slope and peak amplitude. We further used cluster-corrected rmANOVAs over time to establish the effects of target vs. non-target information, spatial attention, cue validity and drug on cue-locked and stimulus-locked ERP data. Lastly, we extracted activity from significant clusters, to test for other main and interaction effects.

We analyzed stimulus-locked ERPs with a cluster-corrected permutation (10000 permutations) 2x2x3 (cue x stimulus identity x drug) rmANOVA. For the cue-locked ERP (**Figure 5A**), we performed a cluster-corrected permutation (10000 permutations) 3x2 (drug x hemisphere) rmANOVA.

#### EEG - time-frequency analysis

To test the effects of spatial attention and drug condition on cortical excitability, we calculated the lateralization of TF power (contralateral – ipsilateral) across cue validity and drug conditions and then tested this lateralization against zero with cluster-corrected two-sided permutation tests (10000 permutations; **Figure 5C**). Furthermore, we extracted power from the alpha-band (8-12Hz) and tested the effects of hemisphere (contralateral vs. ipsilateral to cue) and drug condition on power within this frequency band with a cluster-corrected permutation (10000 permutations) 3x2 (drug x hemisphere) rmANOVA. Lastly, we extracted lateralized single trial alpha power from the late cluster plotted in **Figure 5D** and binned data according to prestimulus alpha power in two evenly sized bins. Then, for every bin and cue validity condition we calculated d’ and tested the effects of drug, prestimulus alpha power bin and cue validity on d’ with a 3x2x2 rmANOVA (**Figure 5 – Supplement 2**).

## Supplementary Figures

**Figure 1 – Supplement 1.**
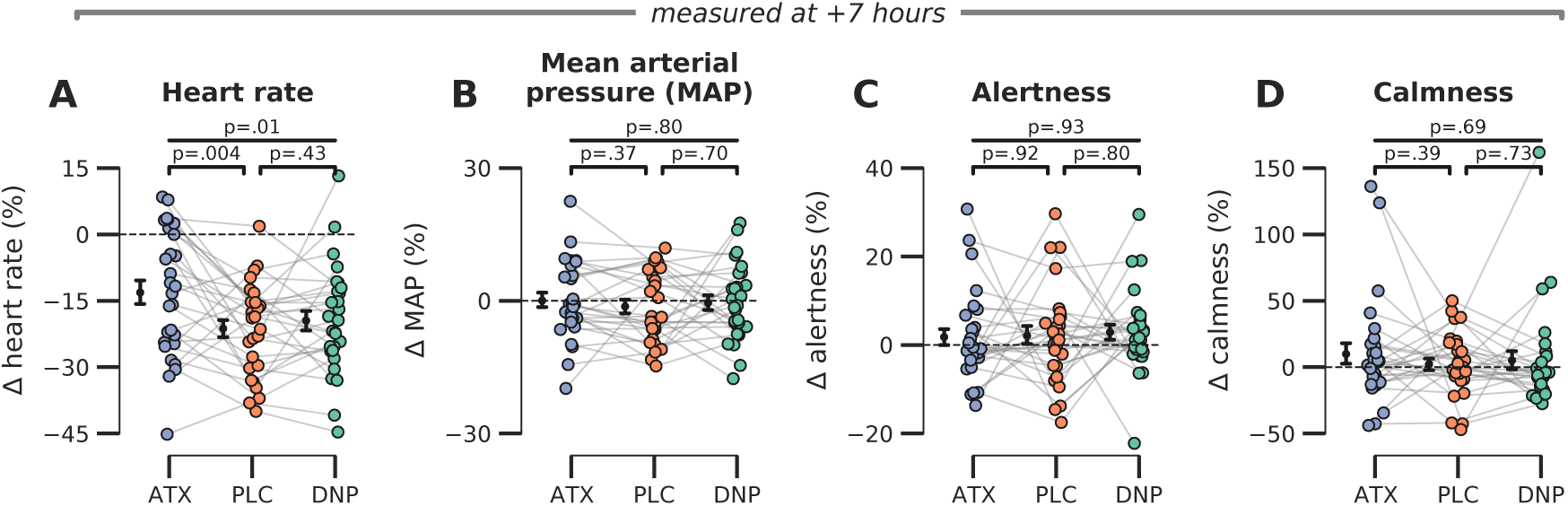
Late effects of drug on subjective and bodily measures of arousal. **A)** Heart rate was modulated by drug condition at the end of the experimental session (F_2,54_=5.44, p=.01, ƞ^2^_*p*_=0.17) and post-hoc tests showed that this effect was driven by increased heart rates under ATX vs. PLC (t(27)=3.11, p=.004, ƞ^2^_*p*_=0.26) but not DNP vs. PLC t(27)=0.80, p=.43, ƞ^2^_*p*_=0.02, BF_01_=3.72). **B-D)** We observed no late effects of drug condition on **B)** mean arterial blood pressure (F_2,54_=0.23, p=.80, ƞ^2^_*p*_=0.01), **C)** subjective ratings of alertness (F_2,54_=0.37, p=.69, ƞ^2^_*p*_=0.01), and **D)** calmness (F_2,54_=0.03, p=.97, ƞ^2^_*p*_=0.00) Error bars around the means indicate SEM. *Abbr.*: ATX: atomoxetine, PLC: placebo, DNP: donepezil.

**Figure 2 – Supplement 1.**
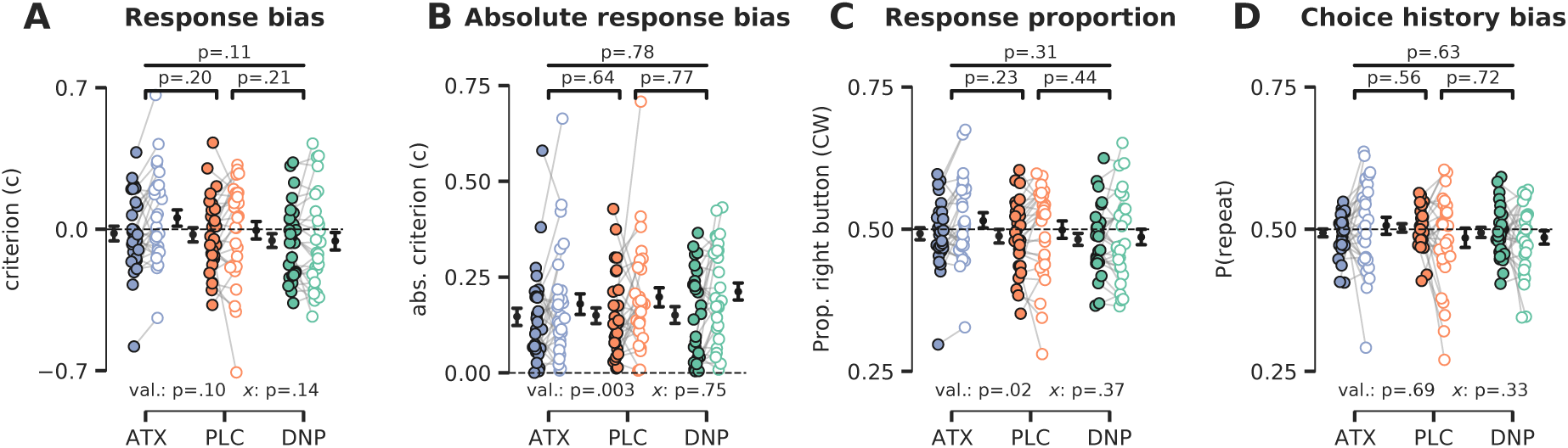
Cue validity and drug condition effects on (choice history) bias. **A)** SDT criterion was not modulated by cue validity (F_1,27_=2.89, p=.10, ƞ^2^_*p*_=0.10) or drug (F_2,54_=2.31, p=.11, ƞ^2^_*p*_=0.08), nor was there a modulatory effect of drug on the effects of cue validity (F_2,54_=2.01, p=.14, ƞ^2^_*p*_=0.07). **B)** Absolute SDT criterion was minimized for validly cued trials, suggesting that participants were more biased (either liberal or conservative) when they did not attend the target stimulus (F_1,27_=11.05, p=.003, ƞ^2^_*p*_=0.29). There were no drug condition main effects (F_2,54_=0.25, p=.78, ƞ^2^_*p*_=0.01) and interaction effects (F_2,54_=0.29, p=.75, ƞ^2^_*p*_=0.01). **C)** Participants pressed the right button more often during invalidly cued trials (F_1,27_=5.79, p=.02, ƞ^2^_*p*_=0.18), possibly related to defaulting back to their preferred hand (right-handedness), but there was no main effect of drug condition (F_2,54_=1.19, p=.31, ƞ^2^_*p*_=0.04) nor an interaction between cue validity and drug (F_2,54_=1.02, p=.37, ƞ^2^_*p*_=0.04). **D)** We did not observe any effects of drug condition (F_2,54_=0.46, p=.63, ƞ^2^_*p*_=0.02), cue validity (F_1,27_=0.17, p=.69, ƞ^2^_*p*_=0.01) or their interaction (F_2,54_=1.14, p=.33, ƞ^2^_*p*_=0.04) on choice history bias. Note that *x* demarks the omnibus interaction between drug condition and cue validity. Val. (short for validity) refers to the factor cue validity.

**Figure 2 – Supplement 2.**
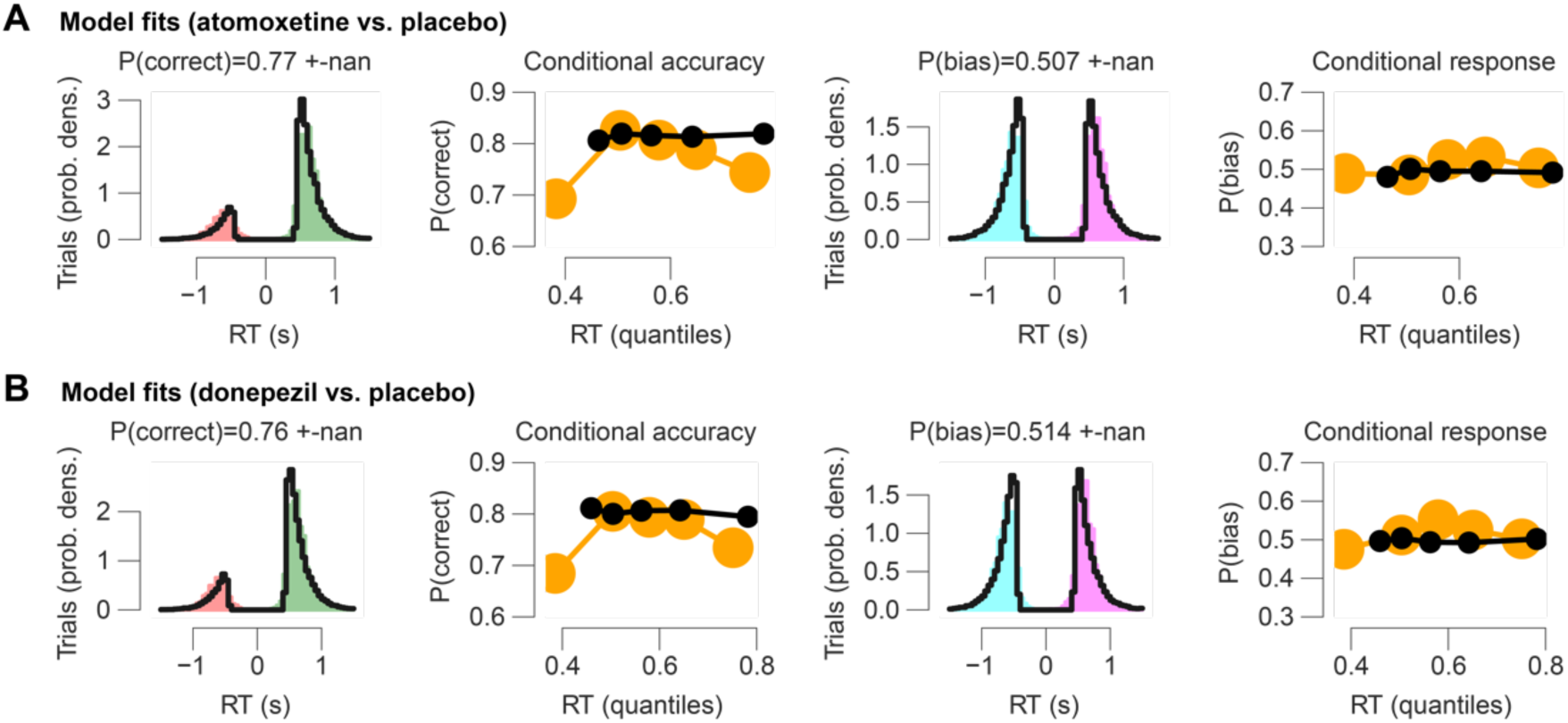
HDDM model fits. All model fits are shown with respect to the empirical data after collapsing across conditions (cue validity and drug). Note that our modeling approach renders the DDM fits complicated to interpret, as the intercept of the effect-coded regression implementation of the DDM reflects a grand mean across conditions. Therefore, model fits show how well empirical data fits the model across all experimental conditions. **A)** Model fits for the atomoxetine (vs. placebo) model. Left panel shows RT fits (black lines) for incorrect and correct responses. Empirical RTs for correct responses are shown in green, for erroneous responses in red. The second panel (from the left) shows modelled proportion correct (black dots) within 5 quantiles of modelled RTs, compared to empirical proportion correct responses (yellow dots) within 5 quantiles of empirical RTs. Right two panels are similar to left two, but show RT distributions for left (turquoise) vs. right (purple) button responses and the proportion of right button responses within each RT bin. **B)** Same as panel **A**, but for the donepezil (vs. placebo) model. The first quintile seems relatively poorly fitted. This is because the model was fitted after excluding outlier RTs (5% of data; **Methods**; Wiecki et al., 2013), which are typically very short-latency errors. Thus, these outlier RTs have affect the first empirical quintile (yellow), but not on synthetic data generated by the fitted model (black).

**Figure 2 – Supplement 3.**
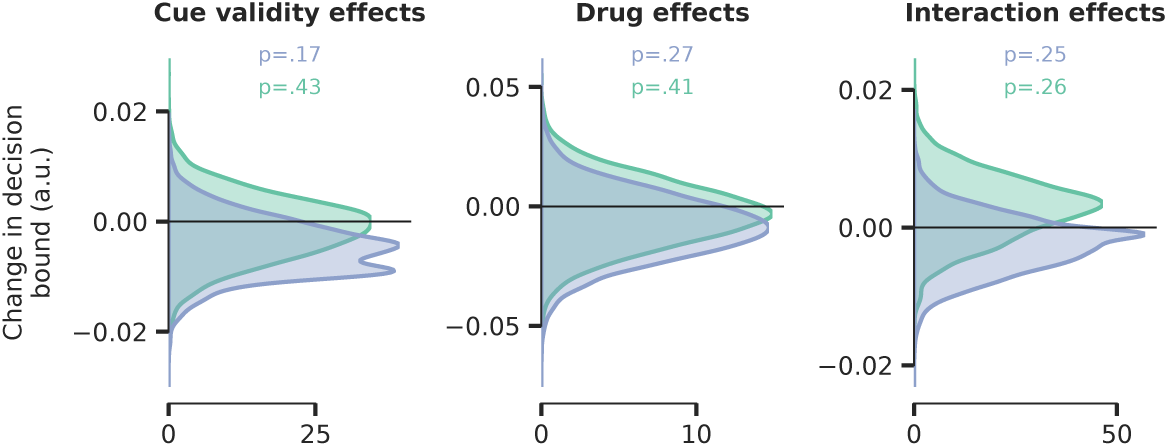
No effects of cue validity, drug, and their interaction on decision bound separation in weighted regression DDM. The computational model described in the main text allowed drift rate, non-decision time and decision bound separation to fluctuate with drug and cue validity. There were no effects of cue validity and drug on decision boundary separation. Blue distribution = ATX model, green distribution = DNP model.

**Figure 2 – Supplement 4.**
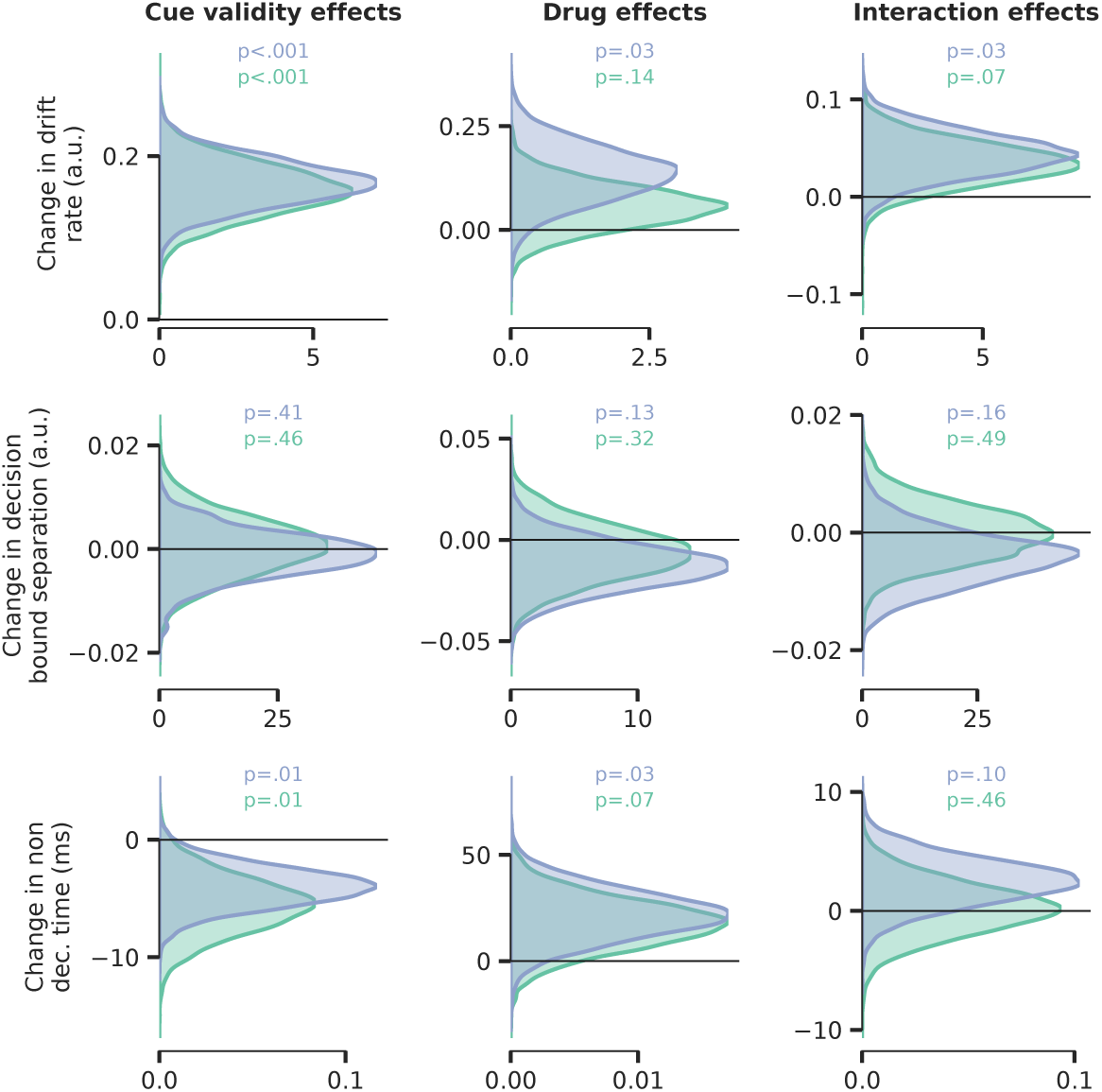
Parameter estimates of unweighted regression DDM. We verified whether applying weights to our effect coding scheme for the regression DDMs, used to counteract the disbalance in the proportion of validly and invalidly cued trials, affected our parameter estimates in any way by fitting the exact same models but now using regular, unweighted, effect coding for cue validity (-1 and 1).There were no substantial differences in parameter estimates compared to the model that used weighted effect coding. Blue distribution = ATX model, green distribution = DNP model.

**Figure 2 – Supplement 5.**
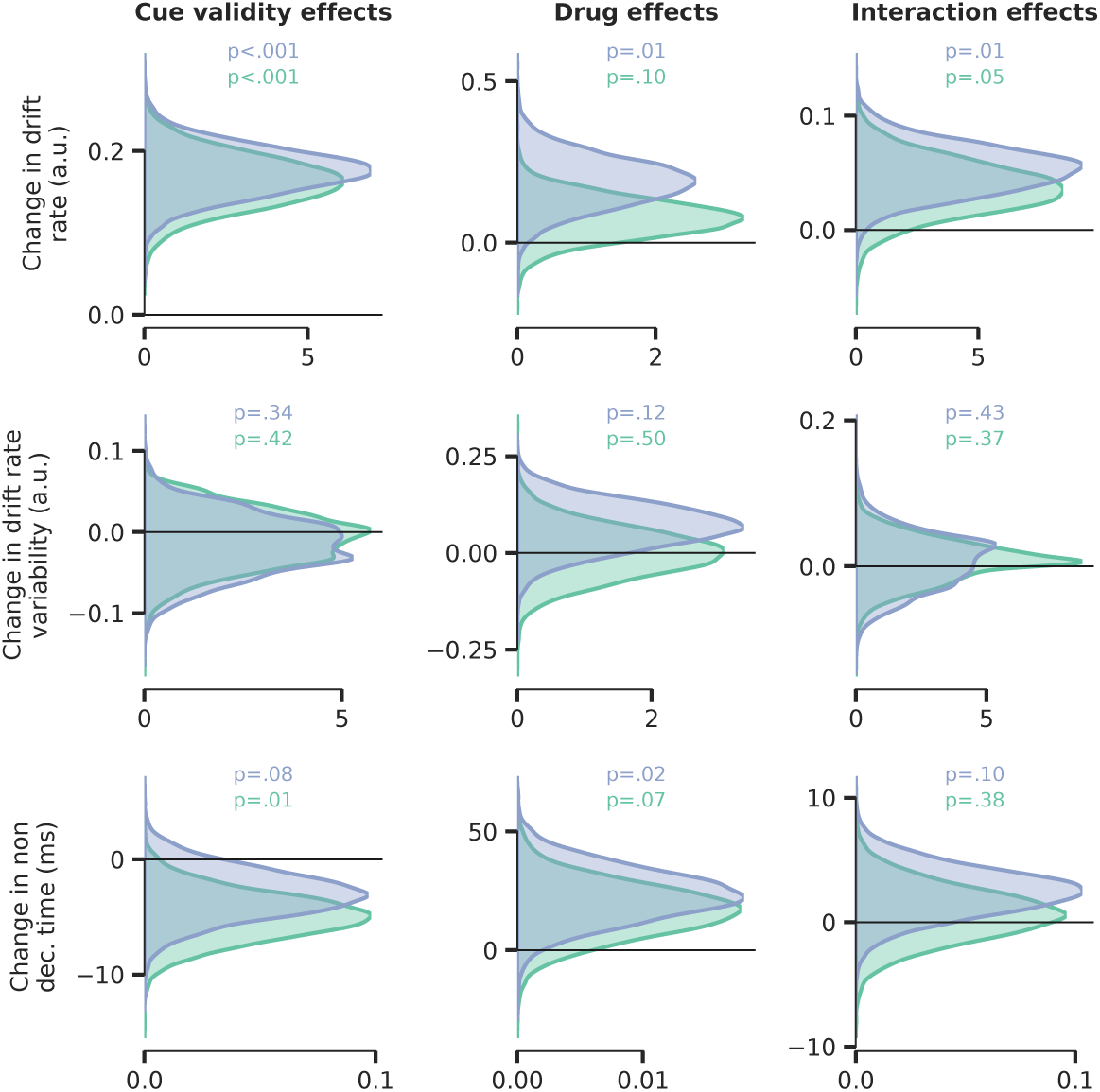
Drift rate variability was not modulated by drug or cue validity. This additional model allowed drift rate (top row), drift rate variability (middle row) and non-decision time (bottom row) to fluctuate with cue validity, drug condition and their interaction, but decision bound separation was fixed across conditions. We observed no significant effects on drift rate variability. Blue distribution = ATX model, green distribution = DNP model.

**Figure 3 – Supplement 1.**
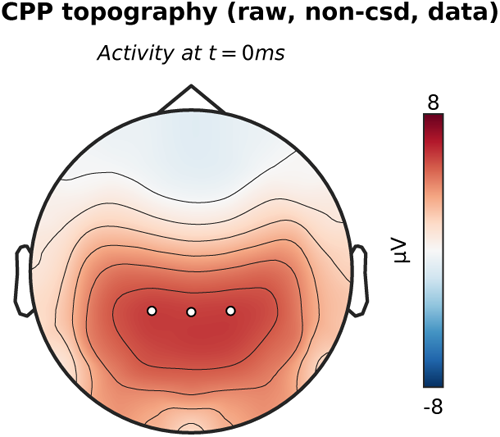
CPP topography without CSD-transformation. The topographic map shows activation at the moment of the response, with white markers indicating the centro-parietal ROI used for the CPP analyses (channels CP1, CP2, CPz).

**Figure 4 – Supplement 1.**
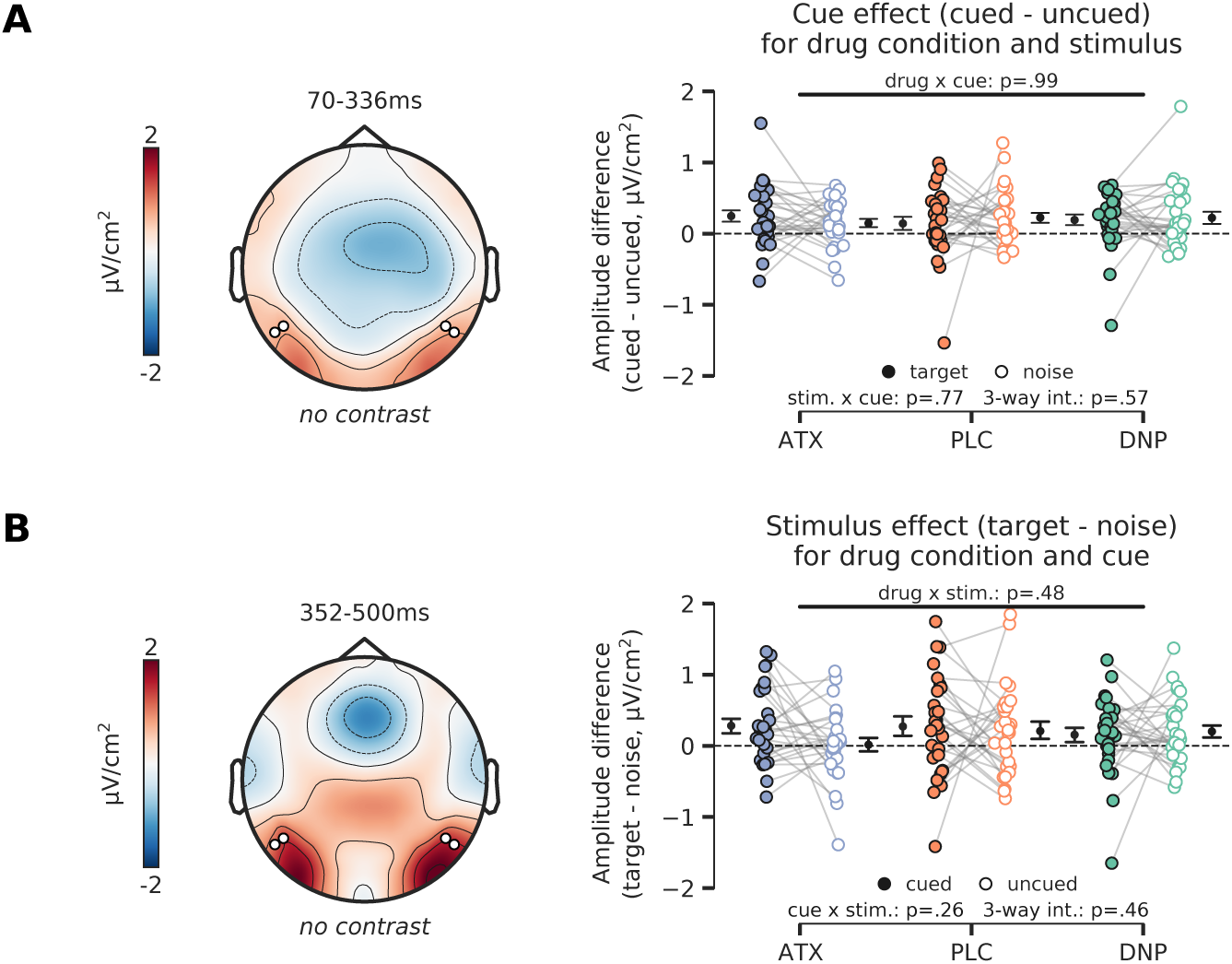
**A)** Topographic map (left panel) shows overall activity (no contrast) across all conditions in the time-window in which we observed a main effect of cue (Figure 4C, left panel). The cue effect (difference between cued and uncued) is plotted in the right panel, separately for targets vs. noise stimuli and for drug condition (ATX/DNP/PLC). Post-hoc tests revealed that the cue effect was not modulated by drug condition, stimulus identity (target or noise), nor was there a three-way interaction between cue, stimulus identity and drug condition. **B)** Same as **A**, but for the stimulus identity effect. The stimulus identity effect (difference between target and noise stimulus) is plotted in the right panel split up for drug and cue (cued/uncued) conditions. The stimulus effect was not modulated by the drug and cue conditions and we also did not observe a three-way interaction effect.

**Figure 5 – Supplement 1.**
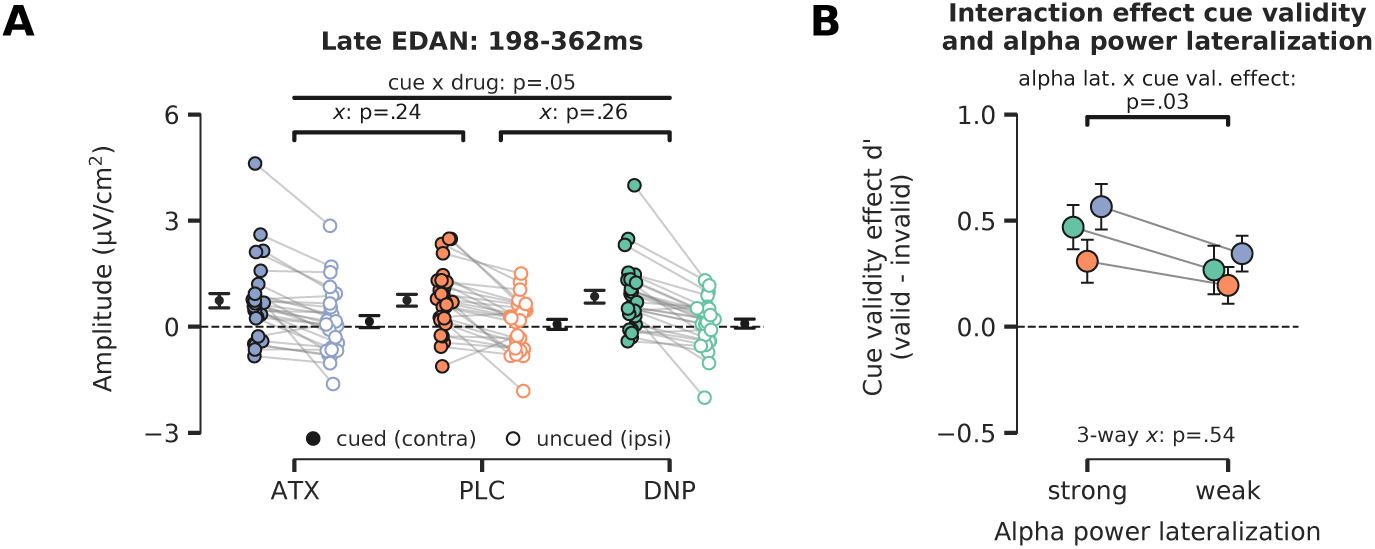
**A)** Late cue-locked activity. The difference in activity between hemispheres associated with the cued and uncued location (contra minus ipsi) for the time window of 198ms-362ms. A trending interaction between cue and drug was observed, but pairwise post-hoc tests showed no reliable differences for ATX/DNP vs. PLC separately. Note that *x* demarks p-value for pairwise (drug vs. PLC) interaction effect. **B)** To test whether alpha-band lateralization bears any relation to task performance, we grouped trials in two bins according to the level of alpha-band suppression in the late time window (median split: strong vs weak alpha-band lateralization). We observed that trials with strong alpha power lateralization were associated with better performance when targets were validly cued vs when they were invalidly cued (F_1,27_=5.14, p=.03, ƞ^2^_*p*_=0.16). Efficient allocation of attention thus aids performance for validly cued trials, but harms performance when the target is presented at the unattended location. Enhanced neuromodulatory activity did not alter this relationship (3-way interaction: F_2,54_=0.63, p=.54, ƞ^2^_*p*_=0.02, BF_01_=4.33).

## References

Alilović, J., Lampers, E., Slagter, H. A., & van Gaal, S. (2023). Illusory object recognition is either perceptual or cognitive in origin depending on decision confidence. PLoS Biology, 21(3), e3002009. 10.1371/journal.pbio.3002009

Alilović, J., Timmermans, B., Reteig, L. C., van Gaal, S., & Slagter, H. A. (2019). No Evidence that Predictions and Attention Modulate the First Feedforward Sweep of Cortical Information Processing. Cerebral Cortex, 29(5), 2261–2278. 10.1093/cercor/bhz038

Aston-Jones, G., & Cohen, J. D. (2005). An Integrative Theory of Locus Coereleus-Norepinephrine function: Adaptive Gain and Optimal Performance. Annual Review of Neuroscience, 28(1). 10.1146/annurev.neuro.28.061604.135709

Bauer, M., Kluge, C., Bach, D., Bradbury, D., Heinze, H. J., Dolan, R. J., & Driver, J. (2012). Cholinergic Enhancement of Visual Attention and Neural Oscillations in the Human Brain. Current Biology, 22(5), 397–402. 10.1016/j.cub.2012.01.022

Baumgartner, H. M., Graulty, C. J., Hillyard, S. A., & Pitts, M. A. (2018). Does spatial attention modulate the earliest component of the visual evoked potential? Cognitive Neuroscience, 9(1–2), 4–19. 10.1080/17588928.2017.1333490

Bentley, P., Driver, J., & Dolan, R. J. (2011). Cholinergic modulation of cognition: Insights from human pharmacological functional neuroimaging. Progress in Neurobiology, 94(4), 360–388. 10.1016/j.pneurobio.2011.06.002

Benwell, C. S. Y., Tagliabue, C. F., Veniero, D., Cecere, R., Savazzi, S., & Thut, G. (2017). Prestimulus EEG Power Predicts Conscious Awareness But Not Objective Visual Performance. Eneuro, 4(6), ENEURO.0182-17.2017. 10.1523/ENEURO.0182-17.2017

Beste, C., Adelhöfer, N., Gohil, K., Passow, S., Roessner, V., & Li, S.-C. (2018). Dopamine Modulates the Efficiency of Sensory Evidence Accumulation During Perceptual Decision Making. International Journal of Neuropsychopharmacology, 21(7), 649–655. 10.1093/ijnp/pyy019

Boehler, C. N., Tsotsos, J. K., Schoenfeld, M. A., Heinze, H.-J., & Hopf, J.-M. (2009). The Center-Surround Profile of the Focus of Attention Arises from Recurrent Processing in Visual Cortex. Cerebral Cortex, 19(4), 982–991. 10.1093/cercor/bhn139

Bond, A., & Lader, M. (1974). The use of analogue scales in rating subjective feelings. British Journal of Medical Psychology, 47(3), 211–218. 10.1111/j.2044-8341.1974.tb02285.x

Boucart, M., Bubbico, G., szaffarczyk, S., Defoort, S., Ponchel, A., Waucquier, N., Deplanque, D., Deguil, J., & Bordet, R. (2015). Donepezil increases contrast sensitivity for the detection of objects in scenes. Behavioural Brain Research, 292, 443–447. 10.1016/j.bbr.2015.06.037

Bruel, B. M., Katopodis, V. G., Vries, R. de, Donner, T. H., McGinley, M. J., & Gee, J. W. de. (2022). Auditory accessory stimulus boosts pupil-linked arousal and reduces choice bias (p. 2022.08.28.505585). bioRxiv. 10.1101/2022.08.28.505585

Busch, N. A., Dubois, J., & VanRullen, R. (2009). The Phase of Ongoing EEG Oscillations Predicts Visual Perception. Journal of Neuroscience, 29(24), 7869–7876. 10.1523/JNEUROSCI.0113-09.2009

Capotosto, P., Babiloni, C., Romani, G. L., & Corbetta, M. (2009). Frontoparietal Cortex Controls Spatial Attention through Modulation of Anticipatory Alpha Rhythms. Journal of Neuroscience, 29(18), 5863–5872. 10.1523/JNEUROSCI.0539-09.2009

Carrasco, M. (2011). Visual attention: The past 25 years. Vision Research, 51(13), 1484–1525. 10.1016/j.visres.2011.04.012

Cohen, J. (1988). Statistical Power Analysis for the Behavioral Sciences (2nd ed.). Routledge. 10.4324/9780203771587

Cohen, M. X. (2014). Analyzing Neural Time Series Data: Theory and Practice. MIT Press.

Cools, R., & D’Esposito, M. (2011). Inverted-U-shaped dopamine actions on human working memory and cognitive control. Biological Psychiatry, 69(12), e113–125. 10.1016/j.biopsych.2011.03.028

Dahl, M. J., Mather, M., Sander, M. C., & Werkle-Bergner, M. (2020). Noradrenergic Responsiveness Supports Selective Attention across the Adult Lifespan. Journal of Neuroscience, 40(22), 4372–4390. 10.1523/JNEUROSCI.0398-19.2020

Dahl, M. J., Mather, M., & Werkle-Bergner, M. (2022). Noradrenergic modulation of rhythmic neural activity shapes selective attention. Trends in Cognitive Sciences, 26(1), 38–52. 10.1016/j.tics.2021.10.009

de Gee, J. W., Colizoli, O., Kloosterman, N. A., Knapen, T., Nieuwenhuis, S., & Donner, T. H. (2017). Dynamic modulation of decision biases by brainstem arousal systems. ELife, 6, e23232. 10.7554/eLife.23232

de Gee, J. W., Correa, C. M. C., Weaver, M., Donner, T. H., & van Gaal, S. (2021). Pupil Dilation and the Slow Wave ERP Reflect Surprise about Choice Outcome Resulting from Intrinsic Variability in Decision Confidence. Cerebral Cortex (New York, N.Y.: 1991), 31(7), 3565–3578. 10.1093/cercor/bhab032

de Gee, J. W., Knapen, T., & Donner, T. H. (2014). Decision-related pupil dilation reflects upcoming choice and individual bias. Proceedings of the National Academy of Sciences, 111(5), E618–E625. 10.1073/pnas.1317557111

de Gee, J. W., Tsetsos, K., Schwabe, L., Urai, A. E., McCormick, D., McGinley, M. J., & Donner, T. H. (2020). Pupil-linked phasic arousal predicts a reduction of choice bias across species and decision domains. ELife, 9, e54014. 10.7554/eLife.54014

Dehaene, S., Changeux, J.-P., & Naccache, L. (2011). The Global Neuronal Workspace Model of Conscious Access: From Neuronal Architectures to Clinical Applications. In Research and Perspectives in Neurosciences (Vol. 18, pp. 55–84). 10.1007/978-3-642-18015-6_4

Desimone, R., & Duncan, J. (1995). Neural mechanisms of selective visual attention. Annual Review of Neuroscience, 18, 193–222. 10.1146/annurev.ne.18.030195.001205

Donohue, S. E., Schoenfeld, M. A., & Hopf, J.-M. (2020). Parallel fast and slow recurrent cortical processing mediates target and distractor selection in visual search. Communications Biology, 3(1), Article 1. 10.1038/s42003-020-01423-0

Forstmann, B. U., Ratcliff, R., & Wagenmakers, E.-J. (2016). Sequential Sampling Models in Cognitive Neuroscience: Advantages, Applications, and Extensions. Annual Review of Psychology, 67, 641–666. 10.1146/annurev-psych-122414-033645

Gelbard-Sagiv, H., Magidov, E., Sharon, H., Hendler, T., & Nir, Y. (2018). Noradrenaline Modulates Visual Perception and Late Visually Evoked Activity. Current Biology, 28(14), 2239–2249.e6. 10.1016/j.cub.2018.05.051

Gold, J. I., & Shadlen, M. N. (2007). The Neural Basis of Decision Making. Annual Review of Neuroscience, 30(1), 535–574. 10.1146/annurev.neuro.29.051605.113038

Gramfort, A., Luessi, M., Larson, E., Engemann, D., Strohmeier, D., Brodbeck, C., Goj, R., Jas, M., Brooks, T., Parkkonen, L., & Hämäläinen, M. (2013). MEG and EEG data analysis with MNE-Python. Frontiers in Neuroscience, 7, 267. 10.3389/fnins.2013.00267

Gratton, C., Yousef, S., Aarts, E., Wallace, D. L., D’Esposito, M., & Silver, M. A. (2017). Cholinergic, But Not Dopaminergic or Noradrenergic, Enhancement Sharpens Visual Spatial Perception in Humans. The Journal of Neuroscience, 37(16), 4405– 4415. 10.1523/JNEUROSCI.2405-16.2017

Green, D. M., & Swets, J. A. (1966). Signal detection theory and psychophysics (pp. xi, 455). John Wiley.

Händel, B. F., Haarmeier, T., & Jensen, O. (2011). Alpha Oscillations Correlate with the Successful Inhibition of Unattended Stimuli. Journal of Cognitive Neuroscience, 23(9), 2494–2502. 10.1162/jocn.2010.21557

Harris, K. D., & Thiele, A. (2011). Cortical state and attention. Nature Reviews Neuroscience, 12(9), Article 9. 10.1038/nrn3084

Hasselmo, M. E., & Sarter, M. (2011). Modes and Models of Forebrain Cholinergic Neuromodulation of Cognition. Neuropsychopharmacology, 36(1), 52–73. 10.1038/npp.2010.104

Hillyard, S. A., & Anllo-Vento, L. (1998). Event-related brain potentials in the study of visual selective attention. Proceedings of the National Academy of Sciences, 95(3), 781–787. 10.1073/pnas.95.3.781

Hillyard, S. A., Vogel, E. K., & Luck, S. J. (1998). Sensory gain control (amplification) as a mechanism of selective attention: Electrophysiological and neuroimaging evidence. Philosophical Transactions of the Royal Society B: Biological Sciences, 353(1373), 1257–1270.

Howe, W. M., Gritton, H. J., Lusk, N. A., Roberts, E. A., Hetrick, V. L., Berke, J. D., & Sarter, M. (2017). Acetylcholine Release in Prefrontal Cortex Promotes Gamma Oscillations and Theta–Gamma Coupling during Cue Detection. The Journal of Neuroscience, 37(12), 3215–3230. 10.1523/JNEUROSCI.2737-16.2017

Iemi, L., & Busch, N. A. (2018). Moment-to-Moment Fluctuations in Neuronal Excitability Bias Subjective Perception Rather than Strategic Decision-Making. Eneuro, 5(3), ENEURO.0430-17.2018. 10.1523/ENEURO.0430-17.2018

Iemi, L., Chaumon, M., Crouzet, S. M., & Busch, N. A. (2017). Spontaneous neural oscillations bias perception by modulating baseline excitability. Journal of Neuroscience, 37(4), 807–819. 10.1523/JNEUROSCI.1432-16.2016

Iemi, L., Gwilliams, L., Samaha, J., Auksztulewicz, R., Cycowicz, Y. M., King, J.-R., Nikulin, V. V., Thesen, T., Doyle, W., Devinsky, O., Schroeder, C. E., Melloni, L., & Haegens, S. (2022). Ongoing neural oscillations influence behavior and sensory representations by suppressing neuronal excitability. NeuroImage, 247, 118746. 10.1016/j.neuroimage.2021.118746

Jas, M., Engemann, D. A., Bekhti, Y., Raimondo, F., & Gramfort, A. (2017). Autoreject: Automated artifact rejection for MEG and EEG data. NeuroImage, 159, 417–429. 10.1016/j.neuroimage.2017.06.030

JASP TEAM. (2022). JASP (0.16.3) [Computer software].

Jensen, O., & Mazaheri, A. (2010). Shaping Functional Architecture by Oscillatory Alpha Activity: Gating by Inhibition. Frontiers in Human Neuroscience, 4, 186. 10.3389/fnhum.2010.00186

Kaernbach, C. (1991). Simple adaptive testing with the weighted up-down method. Perception & Psychophysics, 49(3), 227–229. 10.3758/BF03214307

Kanamori, T., & Mrsic-Flogel, T. D. (2022). Independent response modulation of visual cortical neurons by attentional and behavioral states. Neuron, S0896627322008030. 10.1016/j.neuron.2022.08.028

Kelly, S. P., Lalor, E. C., Reilly, R. B., & Foxe, J. J. (2006). Increases in Alpha Oscillatory Power Reflect an Active Retinotopic Mechanism for Distracter Suppression During Sustained Visuospatial Attention. Journal of Neurophysiology, 95(6), 3844–3851. 10.1152/jn.01234.2005

Kelly, S. P., & O’Connell, R. G. (2013). Internal and External Influences on the Rate of Sensory Evidence Accumulation in the Human Brain. Journal of Neuroscience, 33(50), 19434–19441. 10.1523/JNEUROSCI.3355-13.2013

Kelly, S. P., & O’Connell, R. G. (2015). The neural processes underlying perceptual decision making in humans: Recent progress and future directions. Journal of Physiology-Paris, 109(1), 27–37. 10.1016/j.jphysparis.2014.08.003

Knapen, T., de Gee, J. W., Brascamp, J., Nuiten, S., Hoppenbrouwers, S., & Theeuwes, J. (2016). Cognitive and Ocular Factors Jointly Determine Pupil Responses under Equiluminance. PLOS ONE, 11(5), e0155574. 10.1371/journal.pone.0155574

Lamme, V. A. F., & Roelfsema, P. R. (2000). The distinct modes of vision offered by feedforward and recurrent processing. Trends in Neurosciences, 23(11), 571–579. 10.1016/S0166-2236(00)01657-X

Loughnane, G. M., Brosnan, M. B., Barnes, J. J. M., Dean, A., Nandam, S. L., O’Connell, R. G., & Bellgrove, M. A. (2019). Catecholamine Modulation of Evidence Accumulation during Perceptual Decision Formation: A Randomized Trial. Journal of Cognitive Neuroscience, 31(7), 1044–1053. 10.1162/jocn_a_01393

Loughnane, G. M., Newman, D. P., Bellgrove, M. A., Lalor, E. C., Kelly, S. P., & O’Connell, R. G. (2016). Target Selection Signals Influence Perceptual Decisions by Modulating the Onset and Rate of Evidence Accumulation. Current Biology, 26(4), 496–502. 10.1016/j.cub.2015.12.049

Martinez-Trujillo, J. C., & Treue, S. (2004). Feature-Based Attention Increases the Selectivity of Population Responses in Primate Visual Cortex. Current Biology, 14(9), 744–751. 10.1016/j.cub.2004.04.028

McCormick, D. A. (1989). Cholinergic and noradrenergic modulation of thalamocortical processing. Trends in Neurosciences, 12(6), 215–221. 10.1016/0166-2236(89)90125-2

McGinley, M. J., David, S. V., & McCormick, D. A. (2015). Cortical Membrane Potential Signature of Optimal States for Sensory Signal Detection. Neuron, 87(1), 179–192.

McGinley, M. J., Vinck, M., Reimer, J., Batista-Brito, R., Zagha, E., Cadwell, C. R., Tolias, A. S., Cardin, J. A., & McCormick, D. A. (2015). Waking State: Rapid Variations Modulate Neural and Behavioral Responses. Neuron, 87(6), 1143–1161. 10.1016/j.neuron.2015.09.012

Murphy, P. R., Robertson, I. H., Balsters, J. H., & O’connell, R. G. (2011). Pupillometry and P3 index the locus coeruleus–noradrenergic arousal function in humans. Psychophysiology, 48(11), 1532–1543. 10.1111/j.1469-8986.2011.01226.x

Murphy, P. R., Vandekerckhove, J., & Nieuwenhuis, S. (2014). Pupil-Linked Arousal Determines Variability in Perceptual Decision Making. PLOS Computational Biology, 10(9), e1003854. 10.1371/journal.pcbi.1003854

Murphy, P. R., Wilming, N., Hernandez-Bocanegra, D. C., Prat-Ortega, G., & Donner, T. H. (2021). Adaptive circuit dynamics across human cortex during evidence accumulation in changing environments. Nature Neuroscience, 24(7), 987–997. 10.1038/s41593-021-00839-z

Murray, A. M., Nobre, A. C., & Stokes, M. G. (2011). Markers of preparatory attention predict visual short-term memory performance. Neuropsychologia, 49(6), 1458–1465. 10.1016/j.neuropsychologia.2011.02.016

Newman, D. P., Loughnane, G. M., Kelly, S. P., O’Connell, R. G., & Bellgrove, M. A. (2017). Visuospatial Asymmetries Arise from Differences in the Onset Time of Perceptual Evidence Accumulation. The Journal of Neuroscience, 37(12), 3378–3385. 10.1523/JNEUROSCI.3512-16.2017

Nieuwenhuis, S., Aston-Jones, G., & Cohen, J. D. (2005). Decision making, the P3, and the locus coeruleus—Norepinephrine system. Psychological Bulletin, 131(4), 510–532. 10.1037/0033-2909.131.4.510

Nuiten, S. A., Canales-Johnson, A., Beerendonk, L., Nanuashvili, N., Fahrenfort, J. J., Bekinschtein, T., & Gaal, S. van. (2021, June 14). Preserved sensory processing but hampered conflict detection when stimulus input is task-irrelevant. ELife; eLife Sciences Publications Limited. 10.7554/eLife.64431

O’Connell, R. G., Dockree, P. M., & Kelly, S. P. (2012). A supramodal accumulation-to-bound signal that determines perceptual decisions in humans. Nature Neuroscience, 15(12), 1729–1735. 10.1038/nn.3248

O’Connell, R. G., & Kelly, S. P. (2021). Neurophysiology of Human Perceptual Decision-Making. Annual Review of Neuroscience, 44, 495–516. 10.1146/annurev-neuro-092019-100200

O’Connell, R. G., Shadlen, M. N., Wong-Lin, K., & Kelly, S. P. (2018). Bridging Neural and Computational Viewpoints on Perceptual Decision-Making. Trends in Neurosciences, 41(11), 838–852. 10.1016/j.tins.2018.06.005

Ogura, H., Kosasa, T., Kuriya, Y., & Yamanishi, Y. (2000). Comparison of inhibitory activities of donepezil and other cholinesterase inhibitors on acetylcholinesterase and butyrylcholinesterase in vitro. Methods and Findings in Experimental and Clinical Pharmacology, 22(8), 609–613. 10.1358/mf.2000.22.8.701373

Papaioannou, O., & Luck, S. J. (2020). Effects of eccentricity on the attention-related N2pc component of the event-related potential waveform. Psychophysiology, 57(5). 10.1111/psyp.13532

Parikh, V., Kozak, R., Martinez, V., & Sarter, M. (2007). Prefrontal acetylcholine release controls cue detection on multiple timescales. Neuron, 56(1), 141–154. 10.1016/j.neuron.2007.08.025

Peirce, J. W. (2007). PsychoPy—Psychophysics software in Python. Journal of Neuroscience Methods, 162(1–2), 8–13. 10.1016/j.jneumeth.2006.11.017

Pereira, M., Perrin, D., & Faivre, N. (2022). A leaky evidence accumulation process for perceptual experience. Trends in Cognitive Sciences, 26(6), 451–461. 10.1016/j.tics.2022.03.003

Pfeffer, T., Avramiea, A.-E., Nolte, G., Engel, A. K., Linkenkaer-Hansen, K., & Donner, T. H. (2018). Catecholamines alter the intrinsic variability of cortical population activity and perception. PLOS Biology, 16(2), e2003453. 10.1371/journal.pbio.2003453

Pfeffer, T., Ponce-Alvarez, A., Tsetsos, K., Meindertsma, T., Gahnström, C. J., Brink, R. L. van den, Nolte, G., Engel, A. K., Deco, G., & Donner, T. H. (2021). Circuit mechanisms for the chemical modulation of cortex-wide network interactions and behavioral variability. Science Advances, 7(29), eabf5620. 10.1126/sciadv.abf5620

Pitts, M. A., Metzler, S., & Hillyard, S. A. (2014). Isolating neural correlates of conscious perception from neural correlates of reporting one’s perception. Frontiers in Psychology, 5, 1078. 10.3389/fpsyg.2014.01078

Podvalny, E., King, L. E., & He, B. J. (2021). Spectral signature and behavioral consequence of spontaneous shifts of pupil-linked arousal in human. ELife, 10, e68265. 10.7554/eLife.68265

Posner, M. I. (1980). Orienting of attention. The Quarterly Journal of Experimental Psychology, 32(1), 3–25. 10.1080/00335558008248231

Praamstra, P., & Kourtis, D. (2010). An early parietal ERP component of the frontoparietal system: EDAN not = N2pc. Brain Research, 1317, 203–210. 10.1016/j.brainres.2009.12.090

Ratcliff, R., & McKoon, G. (2008). The Diffusion Decision Model: Theory and Data for Two-Choice Decision Tasks. Neural Computation, 20(4), 873–922. 10.1162/neco.2008.12-06-420

Renart, A., & Machens, C. K. (2014). Variability in neural activity and behavior. Current Opinion in Neurobiology, 25, 211–220. 10.1016/j.conb.2014.02.013

Rogers, S. L., & Friedhoff, L. T. (1998). Pharmacokinetic and pharmacodynamic profile of donepezil HCl following single oral doses. British Journal of Clinical Pharmacology, 46(Suppl 1), 1–6. 10.1046/j.1365-2125.1998.0460s1001.x

Samaha, J., Iemi, L., Haegens, S., & Busch, N. A. (2020). Spontaneous Brain Oscillations and Perceptual Decision-Making. Trends in Cognitive Sciences, 24(8), 639–653. 10.1016/j.tics.2020.05.004

Samaha, J., Iemi, L., & Postle, B. R. (2017). Prestimulus alpha-band power biases visual discrimination confidence, but not accuracy. Consciousness and Cognition, 54, 47–55. 10.1016/j.concog.2017.02.005

Sánchez-Fuenzalida, N., Gaal, S. van, Fleming, S., Haaf, J. M., & Fahrenfort, J. J. (2022). Predictions and rewards affect decision making but not subjective experience. PsyArXiv. 10.31234/osf.io/5v8dh

Seijdel, N., Loke, J., Klundert, R. van de, Meer, M. van der, Quispel, E., Gaal, S. van, Haan, E. H. F. de, & Scholte, H. S. (2021). On the Necessity of Recurrent Processing during Object Recognition: It Depends on the Need for Scene Segmentation. Journal of Neuroscience, 41(29), 6281–6289. 10.1523/JNEUROSCI.2851-20.2021

Silver, M. A., Shenhav, A., & D’Esposito, M. (2008). Cholinergic Enhancement Reduces Spatial Spread of Visual Responses in Human Early Visual Cortex. Neuron, 60(5), 904–914. 10.1016/j.neuron.2008.09.038

Simpson, D., & Plosker, G. L. (2004). Atomoxetine: A review of its use in adults with attention deficit hyperactivity disorder. Drugs, 64(2), 205–222. 10.2165/00003495-200464020-00005

Soma, S., Shimegi, S., Osaki, H., & Sato, H. (2012). Cholinergic modulation of response gain in the primary visual cortex of the macaque. Journal of Neurophysiology, 107(1), 283–291. 10.1152/jn.00330.2011

Summerfield, C., & Egner, T. (2009). Expectation (and attention) in visual cognition. Trends in Cognitive Sciences, 13(9), 403–409. 10.1016/j.tics.2009.06.003

Theeuwes, J. (2010). Top–down and bottom–up control of visual selection. Acta Psychologica, 135(2), 77–99. 10.1016/j.actpsy.2010.02.006

Thiele, A., & Bellgrove, M. A. (2018). Neuromodulation of Attention. Neuron, 97(4), 769–785. 10.1016/j.neuron.2018.01.008

Thut, G. (2006). -Band Electroencephalographic Activity over Occipital Cortex Indexes Visuospatial Attention Bias and Predicts Visual Target Detection. Journal of Neuroscience, 26(37), 9494–9502. 10.1523/JNEUROSCI.0875-06.2006

Tona, K.-D., Murphy, Peter. R., Brown, S. B. R. E., & Nieuwenhuis, S. (2016). The accessory stimulus effect is mediated by phasic arousal: A pupillometry study: Phasic arousal and the AS effect. Psychophysiology, 53(7), 1108–1113. 10.1111/psyp.12653

Twomey, D. M., Murphy, P. R., Kelly, S. P., & O’Connell, R. G. (2015). The classic P300 encodes a build-to-threshold decision variable. European Journal of Neuroscience, 42(1), 1636–1643. 10.1111/ejn.12936

van Kempen, J., Loughnane, G. M., Newman, D. P., Kelly, S. P., Thiele, A., O’Connell, R. G., & Bellgrove, M. A. (2019). Behavioural and neural signatures of perceptual decision-making are modulated by pupil-linked arousal. ELife, 8, e42541. 10.7554/eLife.42541

van Vugt, M. K., Beulen, M. A., & Taatgen, N. A. (2019). Relation between centro-parietal positivity and diffusion model parameters in both perceptual and memory-based decision making. Brain Research, 1715, 1–12. 10.1016/j.brainres.2019.03.008

VanRullen, R., Busch, N., Drewes, J., & Dubois, J. (2011). Ongoing EEG Phase as a Trial-by-Trial Predictor of Perceptual and Attentional Variability. Frontiers in Psychology, 2. https://www.frontiersin.org/article/10.3389/fpsyg.2011.00060

Velzen, J. van, & Eimer, M. (2003). Early posterior ERP components do not reflect the control of attentional shifts toward expected peripheral events. Psychophysiology, 40(5), 827–831. 10.1111/1469-8986.00083

Vinck, M., Batista-Brito, R., Knoblich, U., & Cardin, J. A. (2015). Arousal and locomotion make distinct contributions to cortical activity patterns and visual encoding. Neuron, 86(3), 740–754. 10.1016/j.neuron.2015.03.028

Wang, X.-J. (2008). Decision Making in Recurrent Neuronal Circuits. Neuron, 60(2), 215–234. 10.1016/j.neuron.2008.09.034

Warren, C. M., Eldar, E., Brink, R. L. van den, Tona, K.-D., Wee, N. J. van der, Giltay, E. J., Noorden, M. S. van, Bosch, J. A., Wilson, R. C., Cohen, J. D., & Nieuwenhuis, S. (2016). Catecholamine-Mediated Increases in Gain Enhance the Precision of Cortical Representations. Journal of Neuroscience, 36(21), 5699–5708. 10.1523/JNEUROSCI.3475-15.2016

Waschke, L., Kloosterman, N. A., Obleser, J., & Garrett, D. D. (2021). Behavior needs neural variability. Neuron, 109(5), 751–766. 10.1016/j.neuron.2021.01.023

Waschke, L., Tune, S., & Obleser, J. (2019). *Neural desynchronization and arousal differentially shape brain states for optimal sensory performance* [Preprint]. Neuroscience. 10.1101/582353

Wiecki, T., Sofer, I., & Frank, M. (2013). HDDM: Hierarchical Bayesian estimation of the Drift-Diffusion Model in Python. Frontiers in Neuroinformatics, 7. https://www.frontiersin.org/articles/10.3389/fninf.2013.00014

Wyart, V., & Koechlin, E. (2016). Choice variability and suboptimality in uncertain environments. Current Opinion in Behavioral Sciences, 11, 109–115. 10.1016/j.cobeha.2016.07.003

Zhou, Y. J., Iemi, L., Schoffelen, J.-M., de Lange, F. P., & Haegens, S. (2021). Alpha Oscillations Shape Sensory Representation and Perceptual Sensitivity. The Journal of Neuroscience, 41(46), 9581–9592. 10.1523/JNEUROSCI.1114-21.2021

